# A dedicated brain circuit controls forward walking in *Drosophila*

**DOI:** 10.64898/2026.01.04.697356

**Authors:** Chris J. Dallmann, Fathima Mukthar Iqbal, Sirin Liebscher, Mert Erginkaya, Sander Liessem, Edda L. Sauer, Hannah Soyka, Federico Cascino-Milani, Jens Goldammer, Kei Ito, Jan M. Ache

**Affiliations:** Neurobiology and Genetics, Julius-Maximilians-University of Würzburg, Würzburg, Germany; Institute of Zoology, University of Cologne, Cologne, Germany

## Abstract

Neural circuits in the brain control high-level parameters of movement, including the initiation, speed, and direction of locomotion. How specific cell types are organized into circuits to compute these control signals and enable context-appropriate behavior remains unclear. Here, we identify central brain neurons in *Drosophila* that can initiate walking and exert graded control over walking speed. Connectome analyses position the neurons on top of a layered brain circuit, which recruits a specific population of descending neurons to control forward walking independently of turning and other parameters. As predicted by the connectome, the circuit enforces straight walking when activated unilaterally, and the activity of the central brain neurons represents a high-level walking drive. This drive is suppressed during flight but flexibly integrated with other control signals to enable complex movement sequences during obstacle negotiation. Together, our findings elucidate a dedicated brain circuit that enables efficient computation, selection, and integration of forward-walking signals for context-appropriate motor control.

## Introduction

Walking and other forms of locomotion require precise control over the start, speed, direction, and termination of movement. Across species and forms of locomotion, these control signals originate from circuits in the brain^1–5^, which recruit specific populations of descending neurons (DNs) to activate motor circuits in the spinal cord or invertebrate nerve cord (Fig. 1a).

**Fig. 1:**
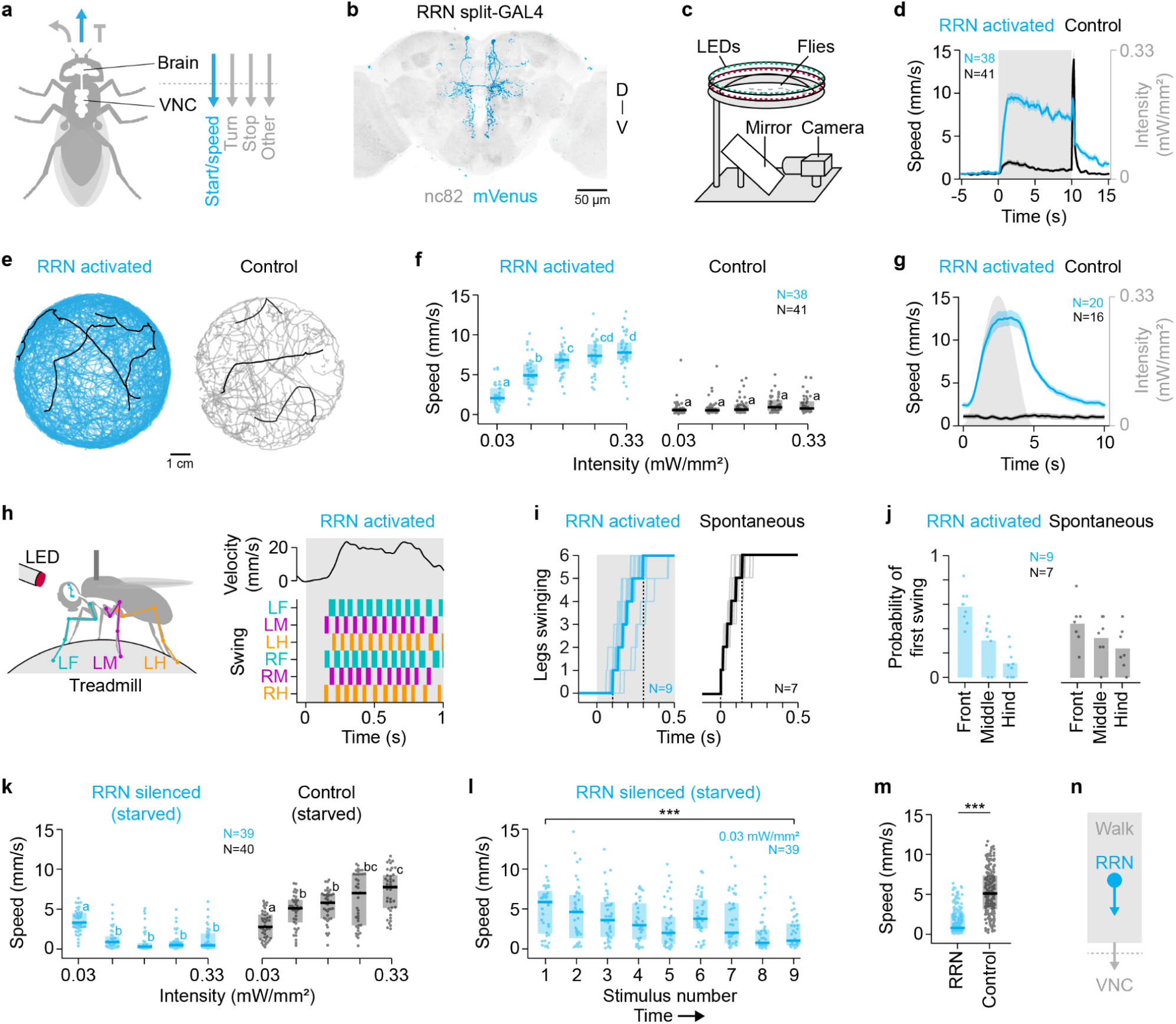
Roadrunner neurons (RRNs) drive walking. **a,** Neural circuits in the *Drosophila* brain control the start and speed of locomotion as well as turning, stopping, and other locomotor parameters. VNC, ventral nerve cord. **b,** Genetic driver line targeting the two RRNs in the central brain. D, dorsal; V, ventral. **c,** Schematic of the free-walking assay. **d,** Translational speed (mean ± s.e.m.) of freely walking RRN>CsChrimson and empty>CsChrimson (control) flies during optogenetic activation (gray). *N* = 38/41 flies (RRN/control); *n* = 9 activations per fly. **e,** Trajectories of flies in the circular arena during optogenetic activation (intensity of 0.33 mW/mm^2^). Black traces, exemplary 10-s trajectories. Sample sizes as in **d**. **f,** Translational speeds of flies during optogenetic activations at different intensities. Dots, animal means; boxes, interquartile range with median; letters, significantly different groups. Sample sizes as in **d**. **g,** Translational speed (mean ± s.e.m.) of flies during sinusoidal optogenetic activation (gray). *N* = 20/18 flies (RRN/control); *n* = 40-50 activations per fly. **h,** Left, Schematic of tethered-walking assay. Right, Leg coordination pattern of a tethered RRN>CsChrimson fly during optogenetic activation. Velocity, forward velocity; bars, swing phases. **i,** Number of legs having started a swing phase relative to the onset of optogenetic activation or spontaneous walking initiation. Thin lines, animal means; thick lines, mean of animal means; black dashed lines, latency to first and last swing phase. *N* = 7/9 flies (RRN/control); *n* = 3-10 transitions per fly. **j,** Probability of a leg to initiate walking. Sample sizes as in **i**. **k,** Translational speed of freely walking RRN>GtACR1 and empty>GtACR1 (control) flies during optogenetic silencing at different intensities. Dots, animal means; boxes, interquartile range with median; letters, significantly different groups. *N* = 39/40 flies (RRN/control); *n* = 9 activations per fly. **l,** Translational speed over consecutive stimulus presentations at the lowest intensity. Inter-stimulus interval is 60 s. Dots, individual animals; boxes, interquartile range with median. *N* = 39 flies. **m,** Translational speed for RRN>GtACR1 and empty>GtACR1 (control) flies, pooled across all stimulus intensities. Dots, animal means for a given intensity. *N* = 39/40 flies (RRN/control); *n* = 5 intensities per fly. **n**, Schematic of RRN within a brain circuit that controls walking (gray box). In **d-g** and **i-m**, data are pooled across females and males.

Recent studies have begun to reveal the organization and function of specific cell types in locomotor circuits in the brain^6–13^. In zebrafish and mice, a subset of glutamatergic neurons in the mesencephalic locomotor region drives specific DNs to control the start and speed of locomotion^6–8,10,14^, while different sets of neurons control turning^9,15–17^. Similarly, in the fruit fly *Drosophila*, central brain neurons and DNs have been identified that specifically control forward locomotion^12,18,19^ or turning^13,20–24^. These studies hint at a potentially shared principle of motor control: the organization of central brain neurons and DNs into parallel functional modules that compute specific, high-level control signals^2^. Parallel modules could enable rapid and reliable computation of locomotor signals and facilitate their context-appropriate selection and integration. However, how specific neurons are organized into such modules is unclear. Moreover, it is unclear how different modules are recruited to form a coherent motor output in the presence of mutually exclusive or synergistic high-level control signals. For example, forward-walking signals should be suppressed during flight but dynamically integrated with turning and stopping signals to enable the complex movement sequences underlying exploration, foraging, courtship, and obstacle negotiation.

In *Drosophila*, we can now directly address these questions by leveraging the recent connectomes of the brain and nerve cord^25–30^ and genetic access to specific cell types^31,32^. Using these tools, recent studies have begun to delineate a circuit that computes turning signals for both walking and flight^13,20–23^. However, the organization and function of circuits that control the initiation and speed of locomotion remains unknown. Neural activity related to forward walking has been measured in several brain regions^33^, but only a few specific central brain neurons and DNs capable of controlling forward walking have been functionally characterized^12,18^. For example, Bolt protocerebral neurons (BPNs) are the only type of central brain neuron known to specifically initiate forward walking and affect walking speed^18^. Owing to the limited number of genetically defined and functionally characterized entry points, the overall organization of walking circuits in the fly brain remains unclear.

Here, we identify and characterize a pair of central brain neurons that can initiate forward walking and modulate walking speed, which we refer to as Roadrunner neurons (RRNs). We leverage brain and nerve cord connectomes to reveal that RRNs and the previously identified BPNs form a layered, forward-walking brain circuit, which recruits a specific population of DNs to drive premotor neurons in the nerve cord. Behavioral experiments confirm that the circuit drives forward walking independently of turning, and in vivo recordings uncover that RRN activity represents a high-level walking drive during spontaneous behavior. Finally, we demonstrate that this walking drive is effectively suppressed during flight but flexibly integrated with turning and climbing signals during obstacle negotiation. Together, our findings reveal how central brain neurons and DNs are organized into a brain circuit dedicated to controlling forward walking. Structural and functional similarities to neurons in the vertebrate brainstem suggest that dedicated locomotor circuits are a shared principle of motor control across species that provides an efficient means for computing, selecting, and integrating high-level control signals for context-appropriate motor control.

### Roadrunner neurons (RRNs) drive walking

To identify walking-related neurons in the central brain of *Drosophila*, we conducted an extensive optogenetic activation screen of sparse genetic driver lines^31^. Activation of one driver line induced particularly fast forward walking. This sparse line drives expression specifically in a pair of central brain neurons (one per hemisphere), without expression in the nerve cord (Fig. 1b and Supplementary Fig. 1a). We refer to these neurons as Roadrunner neurons (RRNs).

We optogenetically activated RRN using CsChrimson^34^ while tracking flies in a circular arena (Fig. 1c; Methods). Bilateral optogenetic activation of RRN caused stationary flies to start walking and walking flies to increase their speed (Fig. 1d). During activation, flies were able to adjust their trajectories similar to spontaneously walking flies. For example, flies turned spontaneously and when encountering other flies or reaching the walls of the arena (Fig. 1e and Supplementary Video 1). Increasing the intensity of optogenetic activation induced progressively higher walking speeds (Fig. 1f; RRN: ANOVA, F(4, 33) = 51.3, p < 0.001; control: ANOVA, F(4, 35) = 1.54, p = 0.19). This intensity-dependent effect was present regardless of the order of the optogenetic stimulus (Supplementary Fig. 1b). When we presented flies with a time-varying, sinusoidal optogenetic stimulus, the speed profile followed the light intensity profile (Fig. 1g). These results demonstrate that RRN is sufficient to initiate walking, and that the combined activity of the left and right RRN can dynamically control walking speed. Females and males showed a similar speed profile (Supplementary Fig. 1c), suggesting that RRN controls walking in general rather than in sex-specific behaviors, such as courtship pursuit.

We next asked whether the leg movements induced by RRN are similar to those of spontaneously walking flies. To test this, we optogenetically activated RRN in flies tethered on an air-supported spherical treadmill and tracked the movements of the legs using two side-view cameras (Fig. 1h, left; Methods). Activation of RRN in stationary flies initiated a normal, coordinated walking pattern (Fig. 1h, right, and Supplementary Video 2). Flies started to swing their legs 103 ± 10 ms (mean ± s.e.m.) after the onset of the optogenetic stimulus and established a coordinated walking pattern about 200 ms later (Fig. 1i). Flies started to move their legs one after the other (Fig. 1i), often but not exclusively starting with a front leg (Fig. 1j). Spontaneously walking flies initiated walking in a similar way (Fig. 1i,j). This suggests that RRN controls walking initiation and speed by activating the endogenous pattern-generating circuits for walking in the nerve cord^5^.

To test whether RRN is required for normal walking, we optogenetically silenced the neurons using GtACR1^35,36^ in freely walking flies. To better assess the silencing effects, we starved flies for 24 h prior to experimentation, which elevates the baseline walking activity^37^. In this assay, control flies walked progressively faster over time, as expected from prolonged starvation (Fig. 1k; ANOVA, F(4, 35) = 18.32, p < 0.001). In contrast, optogenetic silencing of RRN caused flies to slow down over time (Fig. 1k; ANOVA, F(4, 34) = 24.55, p < 0.001). The reduction in speed was already observed over the course of optogenetic stimulus repetitions at the lowest light intensity (Fig. 1l; Mann-Whitney U test, U = 1183, p < 0.001, *N* = 39 flies). Overall, RRN-silenced flies walked significantly slower than control flies (Fig. 1m; Mann-Whitney U test, U = 4787, p < 0.001, *N* = 195-200 silencing periods). These results demonstrate that RRN is required for fast walking.

Together, the optogenetic activation and silencing experiments identify RRNs as critical nodes of a brain circuit that controls walking (Fig. 1n).

### RRNs are part of a forward-walking circuit

To initiate walking and drive changes in walking speed, signals from RRN must travel to motor circuits in the nerve cord via DNs. To investigate the underlying circuit organization, we identified both RRNs in a whole-brain connectome (Fig. 2a, Supplementary Table 1, and Supplementary Video 3; FAFB/FlyWire^26,38^). RRNs are predicted to be cholinergic^39^, suggesting they have an excitatory effect on their postsynaptic partners.

**Fig. 2:**
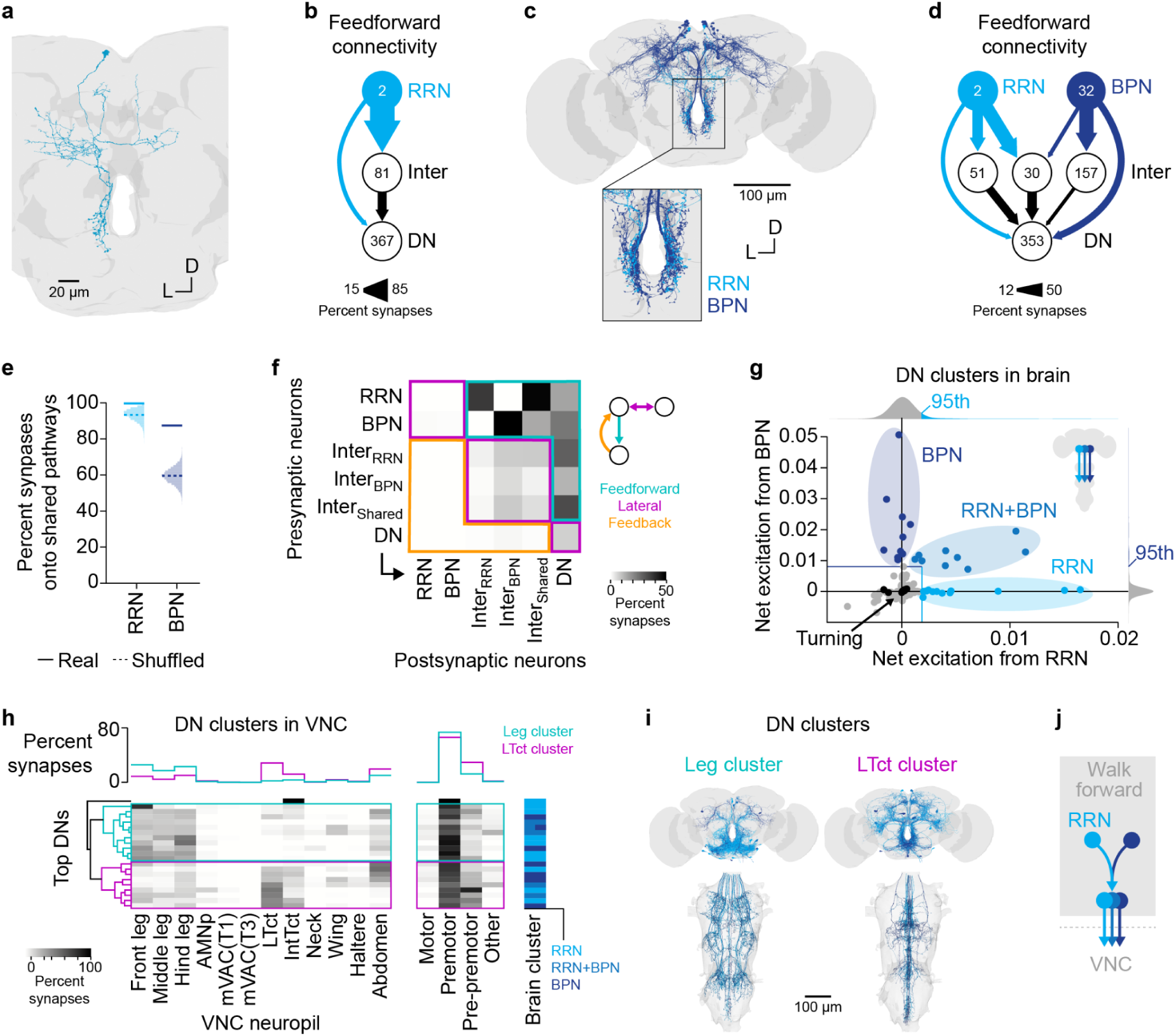
RRNs are part of a forward-walking circuit. **a,** Left RRN in the brain (posterior view; FAFB/FlyWire connectome). D, dorsal; L, left. **b,** Direct and indirect feedforward connectivity from RRN to descending neurons (DNs). Numbers, number of neurons; arrow thickness, percent output synapses. **c,** Overlap of RRN and BPN in the brain. **d,** Feedforward connectivity from RRN and BPN to DNs. The output pathways from BPN onto BPN-specific intermediate neurons and DNs (3% and 4% of output, respectively) is not shown. **e,** Percent synapses from RRN and BPN onto shared DNs, shared intermediate neurons, or intermediate neurons that project to shared DNs. Solid lines, values from the connectome; dashed lines, mean values expected by chance from shuffling connections in the connectome (*n* = 10^4^ shuffles). **f,** Percent synapses from neuron groups of the RRN-BPN circuit in **d** onto each other. Outputs of DNs are relative to their total output in the brain and nerve cord (Methods). **g,** Net excitation from RRN and BPN onto DNs. Clusters, top 5% DNs receiving net excitation from mainly RRN, BPN, or both. Black dots, DNs known to drive turning. Colored lines, 95th percentiles of kernel density estimate (bandwidth = 0.001 net excitation). For DN names, see Supplementary Fig. 2c. **h,** VNC connectivity of DNs highlighted in **g** (MANC connectome). Data for DNs belonging to the same cell type were averaged. For DN names, see Supplementary Fig. 2c. AMNp, accessory mesothoracic neuropil; mVAC, medial ventral association center; T1, prothorax; T3, metathorax; LTct, lower tectulum; IntTct, intermediate tectulum; Brain cluster, cluster assignment of DNs in the brain (from **g**). Note that for one DN type, individual DNs belong to different clusters. **i,** Anatomy of DNs in the brain and VNC, colored by brain cluster and grouped by VNC cluster. **j,** Schematic of convergence of RRN signals onto DNs shared with BPN.

Next, we mapped the mono- and di-synaptic (1- and 2-hop) connections from both RRNs to the layer of DNs (Fig. 2b; Methods). We found that only a small fraction of RRN output (15%) is directly onto DNs. Most output synapses (85%) are formed with intermediate neurons, which in turn have outputs (31%) onto DNs. This suggests that RRNs are higher-level brain neurons that recruit DNs mainly indirectly, forming a layered circuit.

This layered circuit could overlap with circuits that control turning or other locomotor parameters, or it could be dedicated to controlling forward walking. To differentiate between these circuit architectures, we first turned to BPN––the only other known type of excitatory central brain neuron capable of driving forward walking akin to RRN^18^. In contrast to RRNs, BPNs form a large cluster of >30 neurons across the two brain hemispheres (Fig. 2c). The output synapses of RRN and BPN are in similar locations, suggesting that their signals could converge in downstream circuits (Fig. 2c, inset, and Supplementary Video 3). To assess this, we mapped the 1- and 2-hop connections from RRN and BPN onto DNs. As hypothesized, the outputs of RRN and BPN converge at the level of intermediate neurons and DNs (Fig. 2d). Overall, RRN and BPN provide 99.6% and 87.4% of their output onto shared pathways, respectively (Fig. 2d,e; output onto shared DNs, shared intermediate neurons, or intermediate neurons that project to shared DNs). When we shuffled the output connections of each RRN and BPN in this circuit, we found that the observed shared output connectivity was higher than expected by chance (Fig. 2e; Methods).

To better understand the information flow in the shared RRN-BPN circuit, we analyzed the output connectivity of all neurons in the circuit (Fig. 2f). We found that the connections from RRN and BPN to DNs are mainly feedforward (Fig. 2f, teal). This suggests that RRN and BPN are part of a hierarchically-structured circuit in the brain that is dedicated to computing forward-walking control signals.

To further explore this idea, we analyzed how RRN and BPN are positioned to recruit individual DNs. We considered both direct connections and indirect connections via the layer of intermediate neurons, which consists mainly of other central brain neurons with excitatory or inhibitory effects (Supplementary Fig. 2a; Methods). In brief, for each DN targeted by both RRN and BPN, we summed the synaptic input from all excitatory and inhibitory neurons of the shared RRN-BPN circuit relative to its total synaptic input. RRN and BPN reach many (>350) DNs via 2-hop connections, but most of these DNs are predicted to receive very little net excitation or net inhibition (Fig. 2g and Supplementary Fig. 2b). The DNs receiving most net excitation from RRN or BPN (top 5%) fall into three clusters: DNs receiving net excitation mainly from RRN, BPN, or both (Fig. 2g, blue-shaded dots; for cell types, see Supplementary Fig. 2c). The function of these DNs has not been previously investigated. Notably, DNs known to mediate turning signals during walking^13,18,20,21^ receive very little net excitation or net inhibition from RRN and BPN via 2-hop connections (Fig. 2g, black dots, and Supplementary Fig. 2b). Similarly, DNs known to mediate backward walking^40^, stopping^12^, or flight-related signals^19,24^ receive very little net excitation or net inhibition (Supplementary Fig. 2b). The few DNs implicated in initiating forward walking^12,41^ receive comparatively more net excitation (Supplementary Fig. 2b). This connectivity supports the idea that RRNs are part of a forward-walking-specific circuit in the brain, and that turning and other locomotor parameters are controlled by parallel circuits.

In line with this idea, RRN and BPN receive little input from the central complex (CX) and the lateral accessory lobe (LAL)––two brain regions known to compute turning signals^20,22,23^ (Supplementary Fig. 2e). Instead, RRN and BPN receive broad input from several other brain regions, including the ventrolateral neuropils (VLNP), which are associated with visual processing^42^, and the superior neuropils (SNP) and ventromedial neuropils (VMNP), in which neural activity correlated with future changes in walking velocity has been observed^33^. Most input stems from the inferior neuropils (INP), which are not well characterized and have not been previously linked to forward walking. Sensory neurons of any modality are several hops removed (Supplementary Fig. 2f). This input connectivity supports the idea that the forward-walking circuit is recruited to control walking in general rather than in a specific sensory context. RRN and BPN receive different relative inputs from upstream brain neuropils, suggesting they play non-redundant, complementary roles in initiating and controlling forward walking.

The three DN clusters downstream of RRN and BPN could engage overlapping or distinct premotor circuits in the nerve cord. To investigate this, we analyzed the output connectivity of the DNs in a nerve cord connectome (MANC^28,43,44^; Fig. 2h; Methods). One DN cell type (DNp52) has only few synapses in the nerve cord and makes most of its outputs onto central brain neurons. The other DNs have outputs in the leg neuropils (Fig. 2h, left), consistent with the walking-specific effects of RRN and BPN. Based on their projection patterns in the nerve cord, these DNs form two broad clusters: DNs mainly projecting to the leg neuropils (71% of output on average; Fig. 2h, teal), and DNs mainly projecting to the lower tectulum (31% of output; Fig. 2h, magenta), an intermediate nerve cord neuropil thought to be important for coordinating motor control of the legs and wings^45,46^. DNs in both clusters also project to neuropils that control the abdomen and wings. These projection patterns suggest that the DNs control related but distinct functions across the body during walking. Notably, each nerve cord cluster contains DNs from each brain cluster (Fig. 2h, column ‘brain cluster’). This suggests that the excitatory effects of RRN and BPN converge at the DN level and have synergistic effects on motor circuits in the nerve cord.

In the nerve cord, the DNs mainly target a layer of premotor neurons (Fig. 2h, right). Since these neurons are directly presynaptic to motor neurons, they are ideally positioned to transform a descending walking drive into rhythmic leg movements^5,46^. Indeed, a subset of the premotor neurons targeted by the forward-walking circuit is thought to be part of a central pattern generator for walking^41^ (Supplementary Fig. 2d). Another subset is involved in gating proprioceptive feedback from the legs during walking^47^ (Supplementary Fig. 2d). This connectivity suggests that the DNs of the forward-walking circuit activate premotor neurons involved in walking control.

Together, our synaptic connectivity analysis indicates that RRN and BPN are higher-level neurons of a layered, forward-walking-specific circuit in the brain, which converges onto a population of DNs that drives premotor neurons in the nerve cord to elicit and control walking (Fig. 2j and Supplementary Fig. 2g).

### The forward-walking circuit enforces straight walking

The forward-walking circuit provides little excitation to DNs known to drive turning (Fig. 2). However, individual RRNs provide synaptic output only to one side of the brain (mainly in the vest and flange neuropils; Fig. 3a). This raises the possibility that the circuit drives turning if the left and right RRN are recruited asymmetrically during walking. Alternatively, the circuit could be structured so that asymmetric RRN activity is transformed into straight walking without a turning bias.

**Fig. 3:**
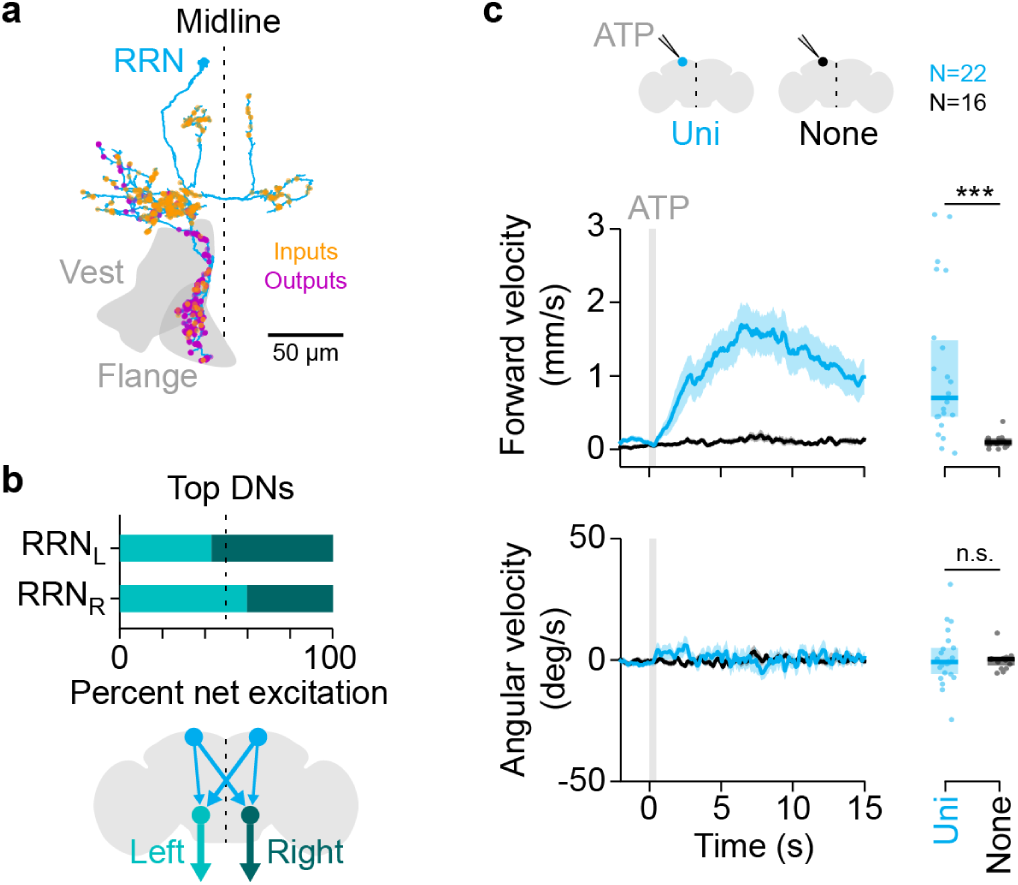
The forward-walking circuit enforces straight walking. **a,** Left RRN with postsynaptic sites (inputs) and presynaptic sites (outputs) in the brain (FAFB/FlyWire connectome). **b,** Percent net excitation of the left and right RRN onto their top 10 DN targets. DNs project to the VNC through the left or right neck connective. Arrows from RRN indicate net excitation across two hops. **c,** Forward and angular velocity (mean ± s.e.m.) of RRN>P2X_2_ (Uni) and RRN>GFP (None, control) flies in response to unilateral application of ATP to a RRN soma. Positive angular velocity corresponds to ipsiversive turning. Dots, animal means (averaged from 0 to 15 s); boxes, interquartile range with median. *N* = 22/16 flies (Uni/None); *n* = 7-54 ATP applications per fly.

When we mapped the 2-hop feedforward connections from the left and the right RRN onto DNs, we found that each neuron provides net excitation onto DNs that project to the left or right side of the nerve cord, in roughly equal proportions (Fig. 3b). This suggests that a unilateral output from RRN is transmitted to both sides of the nerve cord, which should drive forward walking without turning.

To directly test this prediction, we unilaterally activated RRN in tethered flies standing on the spherical treadmill using chemogenetics (Fig. 3c and Supplementary Video 4). We expressed the ATP-receptor P2X_2_ (ref.^48^) in both RRNs and locally applied ATP to either the left or right RRN soma (Methods). Unilateral activation of RRN robustly induced forward walking in every fly tested. The forward velocity of experimental flies increased significantly compared to control flies lacking P2X_2_ expression after ATP application (Fig. 3c, top; Mann-Whitney U test, U = 36, p < 0.001, *N* = 16-22 cells from 16-22 flies). The angular velocity, on the other hand, did not change consistently and remained about zero on average (Fig. 3c, bottom; Mann-Whitney U test, U = 183, p = 0.85, *N* = 16-22 cells from 16-22 flies). That is, flies consistently walked forward but displayed neither an ipsi- nor a contraversive turning bias during unilateral RRN activation.

We confirmed these results by optogenetically activating the left or right RRN in freely walking flies via stochastic expression of CsChrimson using SPARC^32^ (Supplementary Fig. 3a). In line with chemogenetic activation, unilateral optogenetic activation of RRN significantly increased forward walking velocity, with no consistent change in angular velocity (Supplementary Fig. 3b; forward velocity: Mann-Whitney U test, U = 165, p < 0.01; angular velocity; Mann-Whitney U test, U = 101, p = 0.83; *N* = 12-16 flies).

Together, the unilateral activation experiments demonstrate that the forward-walking circuit is structured to transform asymmetric RRN activity into straight walking, confirming the connectome prediction that turning is controlled by circuits running in parallel to the forward-walking circuit. In line with this result, unilateral activation of BPN also drives forward walking without turning^18^.

### RRN activity represents a walking drive

Its activation phenotype and position in the forward-walking circuit predict that RRN encodes a high-level walking drive. If so, RRN activity should be selectively elevated during spontaneous walking and not depend on walking-related sensory feedback.

To test these predictions, we performed whole-cell patch clamp recordings from individual RRNs in tethered walking flies (Fig. 4a and Supplementary Video 5; Methods). As predicted, RRN activity was strongly correlated with walking (Fig. 4a). RRN had a baseline spike rate of about 6 Hz in stationary flies, which significantly increased to about 12 Hz when flies were walking (Fig. 4b; Wilcoxon signed-rank test; W = 0, p < 0.001; *N* = 13 cells from 13 flies). On average, the spike rate and membrane potential increased prior to walking onset (Fig. 4c). Changes in spike rate preceded changes in translational and angular speed by about 100 ms (Fig. 4d; cross correlation, r = 0.61 and r = 0.52, respectively, *N* = 13 cells from 13 flies). When we compared the mean spike rate during slower and faster walking periods for each animal, we found that the spike rate was significantly higher during faster walking (Fig. 4e; 20% lowest speeds versus 20% highest speeds; Wilcoxon signed-rank test; W = 0, p < 0.001, *N* = 13 cells from 13 flies). These results demonstrate that RRN is recruited during spontaneous walking and that increases in RRN activity drive walking, not vice versa.

**Fig. 4:**
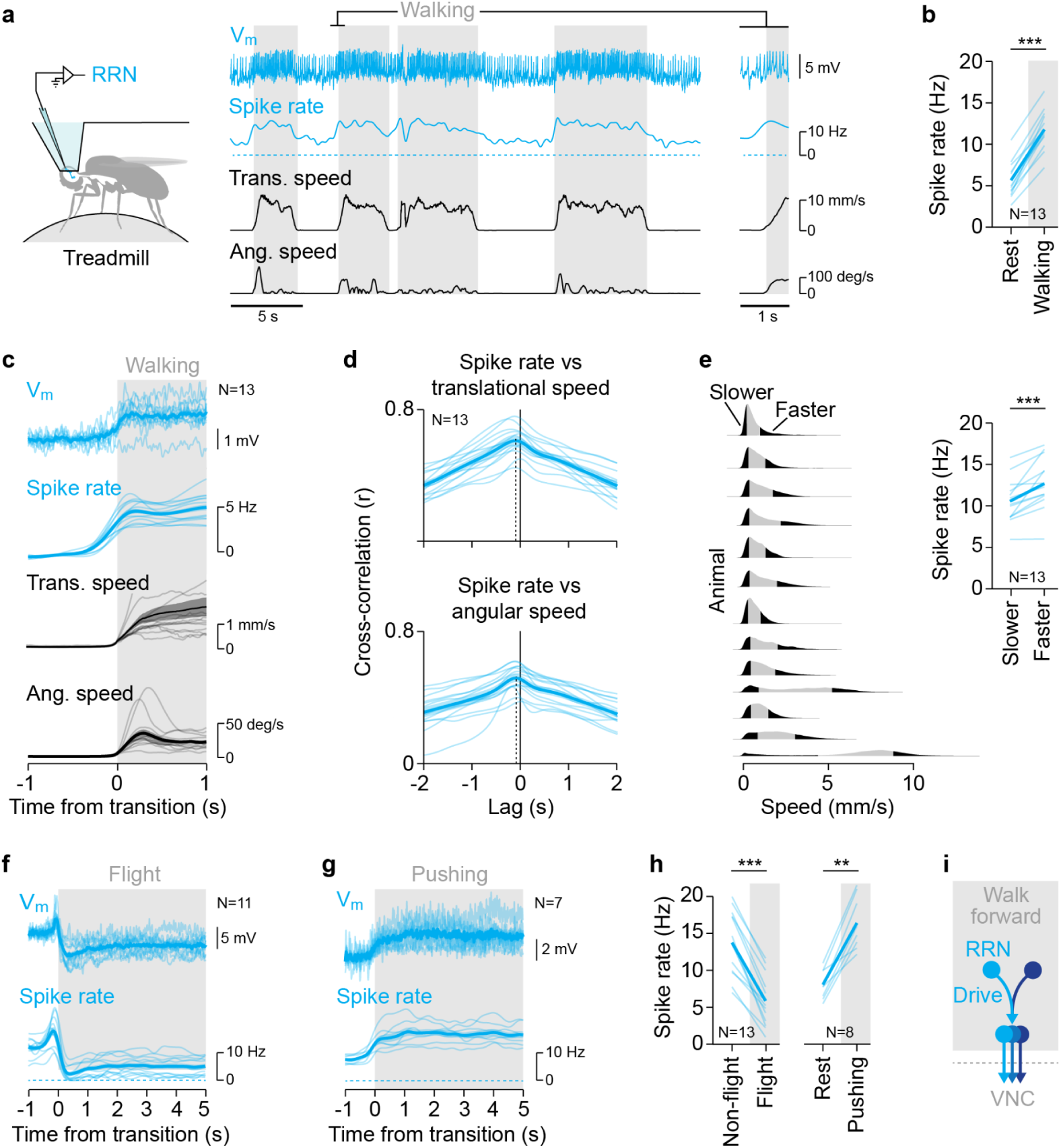
RRN activity represents a walking drive. **a,** Example RRN recording in a tethered walking fly. **b,** Mean spike rate during rest and walking. Thin lines, animal means; thick lines; mean of animal means. *N* = 13 cells from 13 flies. **c,** Mean membrane potential, spike rate, translational speed, and angular speed aligned to the transition into walking. Membrane potential and spike rate are baseline subtracted (mean from -1 to -0.5 s). Membrane potential is smoothed with a 20 ms moving average filter. One speed trace is cut off. Thin lines, animal means; thick lines, mean of animal means; shadings, s.e.m. *N* = 13 cells from 13 flies; *n* = 12-329 transitions per fly. **d,** Cross-correlation between z-scored spike rate and z-scored speed. Dashed lines indicate peak of correlation at around -100 ms. Sample size as in **b**. **e,** Left, Translational speed distributions for each fly tested (kernel density estimate, bandwidth = 0.1 mm s^-1^). Black patches, speeds below the 20th percentile (slower) and above the 80th percentile (faster). Right, Mean spike rate during slower and faster walking. Sample size as in **b**. **f,** Same as **c** but for the transition into flight. Note that the airpuff used to induce flight triggered a few action potentials in a subset of trials. *N* = 11 cells from 11 flies; *n* = 3-10 transitions per fly. **g,** Same as **c** but for the transition into pushing. *N* = 7 cells from 7 flies; *n* = 3-29 transitions per fly. **h,** Left, Mean spike rate during non-flight and flight. *N* = 13 cells from 13 flies. Right, Mean spike rate during rest and pushing. *N* = 8 cells from 8 flies. **i,** Schematic of RRN encoding a walking drive.

To confirm that RRN is specifically active during waking and not generally during locomotion, we recorded RRNs in tethered flying flies (Supplementary Fig. 4a and Supplementary Video 5). The spike rate and membrane potential of RRN decreased strongly at the beginning of flight and remained low throughout flight (Fig. 4f,h; Wilcoxon signed-rank test; spike rate: W = 91; p < 0.001; membrane potential: W = 91; p < 0.001; *N* = 13 cells from 13 flies). The decrease in spike rate and hyperpolarization demonstrate that RRNs are inhibited during flight, confirming that they function specifically in walking control.

Finally, to test whether RRN activity depends on walking-related sensory feedback, we prevented flies from walking by immobilizing the treadmill. In this assay, flies frequently attempted to walk by pushing against the treadmill for extended periods (Supplementary Video 5). During these walking attempts, the spike rate and membrane potential increased significantly (Fig. 4g,h and Supplementary Fig. 4b; Wilcoxon signed-rank test, W = 0, p = 0.016, *N* = 7 cells from 7 flies). Spike rates even exceeded those observed during actual walking (walking: 11.8 ± 0.7 Hz; walking attempts: 16.4 ± 1.3 Hz; Mann-Whitney U test, U = 13, p < 0.01, *N* = 8-13 flies). These results suggest that RRN activity is not driven by walking-related sensory feedback signals from ascending pathways^49,50^ but instead represents an internal walking drive, which is elevated when it cannot be translated into actual walking output.

Together, the in vivo dynamics confirm the connectome prediction that RRN is a high-level, core node of a layered circuit dedicated to controlling spontaneous forward walking (Fig. 4i).

### Walking drive is flexibly suppressed or integrated

In the absence of competing high-level control signals, activation of RRN robustly initiated forward walking (Figs. 1 and 3). However, during natural behavior, competing control signals may exist, such as a drive to groom. To form a coherent motor output, mutually exclusive control signals should be suppressed, whereas synergistic control signals should be integrated. These interactions are difficult to infer from static connectomes and require neural manipulations in different behavioral contexts.

To test how the forward-walking drive interacts with mutually exclusive control signals, we optogenetically activated RRN in grooming or flying flies. When flies were grooming, activation of RRN consistently initiated walking (Fig. 5a and Supplementary Video 6). The transition from grooming to walking was as robust as the transition from standing to walking, although it occurred later on average (Fig. 5a). In contrast, activation of RRN did not terminate flight, nor did it induce leg movements during flight (Fig. 5a and Supplementary Video 6). These results demonstrate that the forward-walking drive is effectively suppressed during flight, in line with our finding that RRN itself is inhibited during flight (Fig. 4). Moreover, the results reveal an action selection hierarchy among mutually exclusive motor behaviors, in which flight overrides walking and walking overrides grooming.

**Fig. 5:**
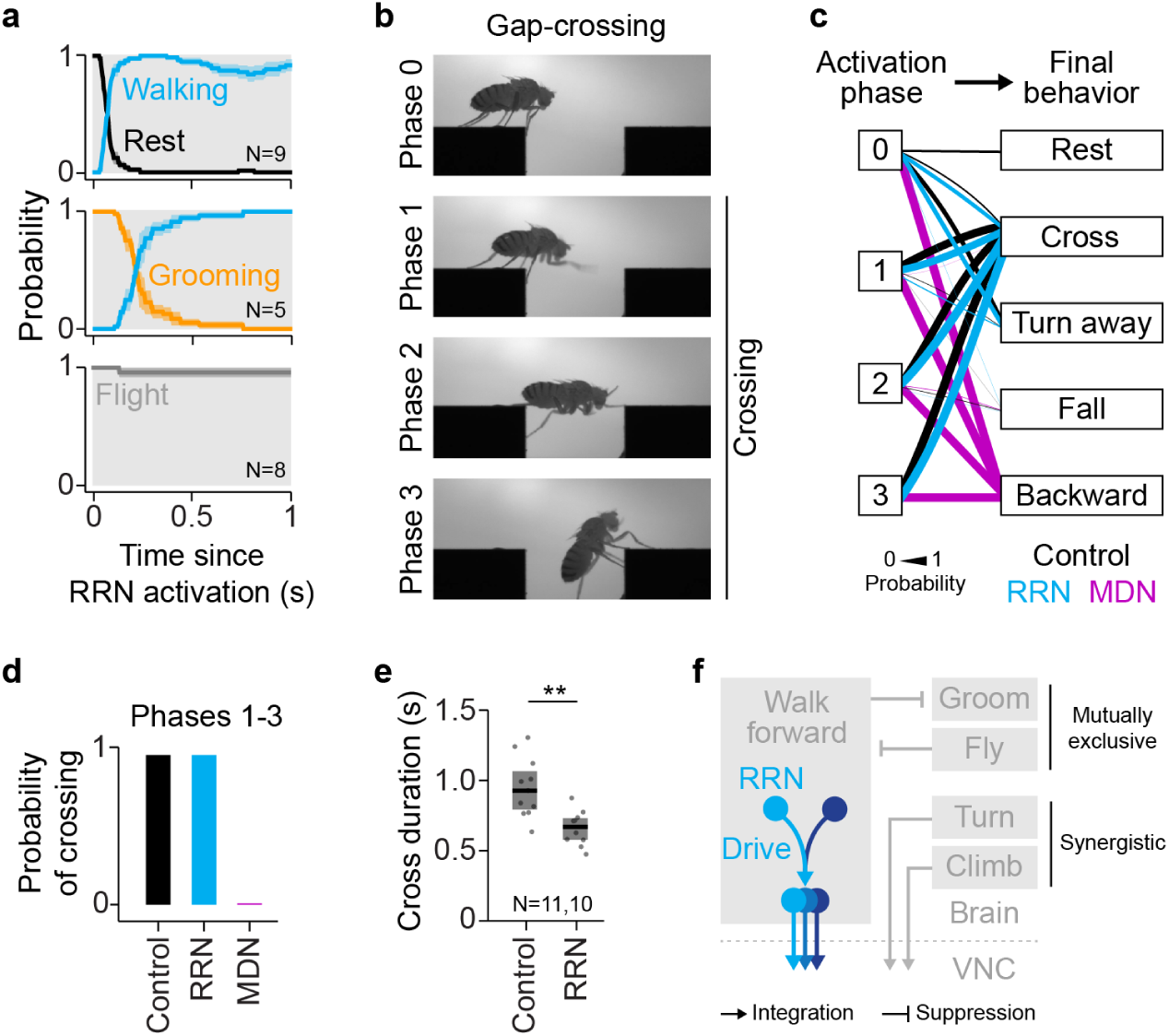
Walking drive is flexibly suppressed or integrated. **a,** Probability of behaviors (mean ± s.e.m.) of tethered RRN>CsChrimson flies during optogenetic activation (gray). *N* = 9/5/8 flies (rest/grooming/flight); *n* = 3-16 activations per fly. **b,** Phases of gap crossing: front leg searching (phase 1), front leg reaching and other legs following (phase 2), walking out of gap (phase 3). Gap width is 3.5 mm. The wings were clipped to prevent flight. **c**, Probability of final behaviors of empty>CsChrimson (control), RRN>CsChrimson, and MDN>CsChrimson flies following optogenetic activation before (phase 0) or during (phases 1-3) gap-crossing. Behaviors pooled across flies. *N* = 7-12/9-11/4-13 flies (control/RRN/MDN). **d,** Probability of successful gap crossing of empty>CsChrimson (control), RRN>CsChrimson, and MDN>CsChrimson flies with optogenetic stimulation during crossing (mean of phases 1-3, see **c**). **e,** Gap-crossing duration for empty>CsChrimson (control) and RRN>CsChrimson flies. Dots, animal means; boxes, interquartile range with median. *N* = 11/10 flies (control/RRN); *n* = 1-10 crosses per fly. **f,** Schematic of suppression of mutually exclusive control signals and integration of synergistic control signals.

To test whether the forward-walking drive is flexibly integrated with synergistic walking control signals, we turned to an assay in which flies cross a water-filled gap^51^ (Fig. 5b). Gap-crossing is a complex obstacle negotiation task, which flies readily perform in a predictable movement sequence^51^ (Fig. 5b). After stepping into the gap, flies briefly stop (phase 0). If flies decide to cross the gap, they initiate climbing by performing search movements with their front legs to locate the opposite side of the gap (phase 1). Once the front legs reach the opposite side, flies climb across the gap with their middle and hind legs (phase 2). Finally, flies walk out of the gap (phase 3). In this motor task, forward-walking signals must be flexibly integrated with synergistic climbing signals (if flies decide to cross the gap) or turning signals (if flies decide to walk away from the gap). Without flexible integration, the forward-walking signals would cause flies to walk straight into the water-filled gap.

We optogenetically activated RRN before and during the different phases of gap-crossing. Optogenetic activation of RRN before gap-crossing always initiated movement (Fig. 5c and Supplementary Fig. 5a, phase 0). But similar to control flies, RRN-activated flies either turned away from the gap or climbed across it, displaying the typical gap-crossing behavior (Fig. 5c, Supplementary Fig. 5a, and Supplementary Video 7). RRN-activated flies also climbed across the gap when the activation occurred during the different phases of gap-crossing, and they crossed the gap as successfully as control flies (Fig. 5c,d and Supplementary Fig. 5a, phases 1-3). At no point did activation of RRN induce ‘simple’ forward walking that drove flies into the water-filled gap. Notably, RRN-activated flies crossed the gap faster than control flies (Fig. 5e; Mann-Whitney U test, U = 99, p < 0.01, *N* = 10 RRN flies, *N* = 11 control flies). We can exclude differences in body size as a trivial explanation for faster crossing, because RRN-activated flies were not significantly larger than control flies (Supplementary Fig. 5b; Mann-Whitney U test, U = 33, p = 0.13, *N* = 10 RRN flies, *N* = 11 control flies). These results demonstrate that the forward-walking control signals are flexibly integrated with climbing and turning signals to support the animal’s behavioral goal during obstacle negotiation.

To confirm that high-level walking signals can interrupt gap-crossing if they are not synergistic, we optogenetically activated Moonwalker descending neurons (MDN or DNp50)—a set of command-like DNs that drive backward walking^40,52^. In line with its direct connectivity to premotor neurons in the nerve cord^55^, activation of MDN reliably elicited movement in the gap-crossing assay, with shorter latencies than activation of RRN (Supplementary Fig. 5c; Mann-Whitney U test, U = 2995.5, p < 0.001, *N* = 9-13 animals). Strikingly, MDN-activated flies always walked backward, rendering flies unable to cross the gap (Fig. 5c,d and Supplementary Fig. 5a). In the last phase of gap-crossing, activation even caused flies to walk backward into the water-filled gap (Supplementary Video 7). These results demonstrate that walking signals can interrupt gap-crossing if they are not synergistic.

Together, the activation experiments demonstrate that the forward-walking control signals are flexibly suppressed or integrated in a context-appropriate manner (Fig. 5f). Suppression prevents interference between mutually exclusive motor behaviors, while integration enables complex movement sequences.

## Discussion

In this study, we uncovered the organization and function of a brain circuit dedicated to controlling forward walking in *Drosophila*. Previous studies had characterized individual central brain neurons and DNs involved specifically in forward walking^12,18^. However, the organization and function of a circuit dedicated to controlling the initiation and speed of forward walking and its interaction with competing locomotor control signals had remained unknown. We identified and characterized Roadrunner neurons (RRNs), a pair of central brain neurons that can initiate walking and modulate walking speed (Fig. 1). This novel genetic and anatomical entry point enabled us to delineate a brain circuit that controls forward walking independently of turning or other locomotor parameters (Figs. 2 and 3). The activity of RRN represents a high-level walking drive (Fig. 4), which is flexibly suppressed or integrated in a context-appropriate manner (Fig. 5).

Our conclusion that RRN is part of a dedicated forward-walking circuit is based on three lines of evidence. First, the connectome revealed that the outputs of RRN converge with those of BPN, the only other type of central brain neuron specifically implicated in driving forward walking in *Drosophila*^18^. RRN and BPN are predicted to have little excitatory impact onto DNs known to drive turning^13,18,20,21^ or other locomotor parameters^19,24,40^. Moreover, RRN and BPN receive little input from the central complex and the lateral accessory lobe, which mediate turning signals^13,22,23^. Second, unilateral activation of RRN resulted in forward walking with no turning bias, as was also shown for BPN^18^. Third, RRN was recruited during spontaneous walking but inhibited during flight. These findings demonstrate that RRN is a higher-level node of a brain circuit dedicated to computing forward-walking control signals.

The boundaries of the forward-walking circuit remain to be explored, but our connectome analysis makes specific predictions that can be tested experimentally in the future. For example, several intermediate neurons receive shared input from RRN and BPN, suggesting they form core nodes in the circuit. Moreover, a population of DNs is predicted to send forward-walking signals to motor circuits in the nerve cord, perhaps to promote flexibility and robust encoding^17^. Our finding that RRN and BPN provide outputs in overlapping brain neuropils, including in the vest and flange, and receive most of their inputs from inferior neuropils, including the inferior clamp, position these neuropils as important hubs for walking control. The broad inputs to RRN and BPN are in line with the idea that the forward-walking circuit is recruited for walking in general rather than in specific contexts. Finally, while there are several similarities between RRN and BPN, there are also key differences. Notably, RRNs are a single pair of neurons in the fly brain, whereas BPNs form a cluster of >30 neurons. This suggests that individual RRNs could have a stronger functional impact than individual BPNs, or that there is functional heterogeneity within the BPN cluster.

Strikingly, the forward-walking circuit shares key characteristics with locomotor circuits in the vertebrate brainstem. Like RRN and BPN, neurons in the mesencephalic locomotor region (MLR) exert graded control over locomotor speed^6,8,53,54^, transform unilateral activation into a bilateral locomotor output^6–8,10,53–55^, and target a population of DNs that is distinct from DNs that mediate turning signals^10,14,16,17,56^. Moreover, different sets of MLR neurons and downstream DNs encode locomotor speed in complementary ways^6,8,16^. Similarly, we found that the walking drive provided by RRN is only moderately elevated during faster walking, suggesting that overall walking speed is encoded by the combined activity of neurons in the forward-walking circuit rather than a single cell type. Such a circuit-level encoding of speed could improve robustness and enable finer-scale control than could be realized by single neurons with limited dynamic range. Overall, the structural and functional similarities with neurons in the brainstem of zebrafish, mice, and other vertebrates suggest that the organization of central brain neurons and DNs into functional modules computing specific control signals is a shared principle across species. By leveraging genetic access to specific cell types and connectomes covering the brain and nerve cord in *Drosophila*, our study details the organization and function of such a circuit.

It is not obvious that locomotion should be controlled by neurons organized into dedicated parallel circuits. A single circuit can, in principle, encode multiple parameters simultaneously through multiplexing, providing an economical solution for numerically small nervous systems^57^. On the other hand, dedicated circuits have several functional advantages. Computing specific locomotor control signals in parallel is likely faster and more reliable than multiplexing the signals. Moreover, dedicated locomotor circuits could facilitate the suppression of mutually exclusive control signals. We found that RRN is inhibited during flight, and because optogenetic activation of RRN did not drive leg movements during flight, neurons downstream of RRN are likely inhibited as well. The suppression is presumably specific to the circuits that control walking, because the legs can be moved during flight for steering and landing^58–61^. The suppression could either originate from flight circuits in the brain^24,62^, or ascending feedback pathways that convey a flight state from the nerve cord^63^. In addition, suppression could emerge from state-dependent neuromodulation, as has been observed in DNs^60^. Conversely, to override grooming, the forward-walking circuit could inhibit grooming circuits in the brain and nerve cord^64–67^.

Dedicated locomotor circuits in the brain could also facilitate the decoding and integration of control signals in downstream motor circuits. We found that the forward-walking drive of RRN is flexibly integrated with synergistic climbing or turning signals to support the animal’s decision to cross a gap or turn away from it. In the free-walking assay, the forward-walking drive was also flexibly integrated with turning signals when flies encountered other flies or reached the walls of the arena. Climbing and turning signals are controlled by brain circuits distinct from the forward-walking circuit: climbing is not displayed during normal walking and can be manipulated independently of walking^51^, and both climbing and turning rely in part on the central complex^13,68^. We propose that the integration of synergistic control signals occurs in shared premotor circuits in the nerve cord, as has been proposed for the spinal cord^2^. This is consistent with a growing body of work suggesting that populations of DNs mediate specific motor control signals — or motor primitives^69^ — which can be flexibly combined to enable complex movement sequences^16,17,21,29,61,70,71^. Our results expand on this idea by revealing how central brain neurons and DNs form a dedicated circuit for computing forward-walking control signals.

Overall, our findings suggest that dedicated locomotor circuits in the brain provide an efficient means for computing, selecting, and integrating high-level control signals for flexible, context-appropriate motor control.

## Methods

### Experimental animals

We used *Drosophila melanogaster* raised on standard cornmeal and molasses medium at 25 °C and 60% humidity on a 12:12-h light:dark cycle. Chemogenetic and electrophysiology experiments were done with females 2-5 days old. Optogenetic experiments were done with females and males 3-6 days old that were kept in the dark on fly food coated with 300 µM all-trans-retinal (R2500, Sigma-Aldrich, Steinheim, Germany) for 2-5 days before the experiment. The genotypes are listed in Supplementary Table 2. The sample sizes are listed in Supplementary Table 3.

### Optogenetic experiments with unrestrained animals

We used a previously described setup for optogenetic activation and silencing^37,72^. In brief, the setup consisted of a 10-cm diameter circular glass arena covered with a watch glass, in which flies could walk freely while being filmed from below with a camera at 20 Hz (Basler acA1300-200um, Basler AG). The arena was illuminated from above by white background light. A ring of 60 LEDs provided infrared light (870 nm) for movement tracking. Two additional rings of LEDs provided red light (625 nm) or green light (525 nm) for CsChrimson or GtACR activation, respectively. The setup was controlled with custom Matlab code.

Cold-anesthetized flies were transferred to the arena and given 10 min to acclimate. In the square-wave optogenetic experiments, 18-22 flies were placed in the arena at a time (10-13 females, 7-10 males). Then, the red or green LEDs were activated at one of five intensities (0.03, 0.11, 0.18, 0.26, 0.33 mW/mm^2^). For each intensity, we recorded a separate video, in which we delivered nine 10-s stimuli with a 60-s inter-stimulus-interval. Videos were recorded 5 min apart. In the SPARC experiments, single flies were placed in the arena and tested at the highest intensity. In the sinusoidal optogenetic activation experiments, we also used single flies. The red LEDs were activated in a sinusoidal pattern ranging from 0 to 0.33 mW/mm^2^. For each fly, we recorded 5 trials, in which we delivered ten 5-s stimuli with a 5-s inter-stimulus-interval. Trials were recorded 2 min apart. Because only single flies were recorded in each trial, fly identity could be tracked across trials.

Each fly’s center of mass and body axis were tracked offline using TRex^73^. The automated tracking was reviewed and identity switches between animals were corrected manually within videos. Walking velocities and speeds were calculated as described in ref.^52^. The tracking data were analyzed with custom Python code.

### Immunohistochemistry

To visualize RRNs, flies were dissected for immunostaining as in ref.^52^. The primary antibodies used were rabbit anti-RFP (1:1000 concentration) and mouse anti-nc82 (for neuropil staining, 1:500 concentration). The secondary antibodies used were goat anti-rabbit Alexa Fluor 555 (1:200 concentration) and goat anti-mouse Alexa Fluor 635 (1:400 concentration). Z-stacks of the brain were acquired using a confocal microscope (Leica TCS SP8) and processed in Fiji^74^.

### Optogenetic experiments with tethered animals

Cold-anesthetized flies were tethered to a 30-gauge needle at the dorsal thorax using ultraviolet-curing glue and then placed on an air-supported ball (polypropylene, 6 mm diameter), which functioned as a spherical treadmill. The treadmill was illuminated by two infrared LEDs (980 nm) and tracked at 50 Hz with two orthogonally placed custom motion sensors^52^. The fly was illuminated by two infrared LEDs (850 nm) from the front and the back. Movements of the legs were recorded at 300 Hz with two side-view cameras (Basler acA1300-200 um, Basler AG) equipped with a zoom lens (Ricoh FL-CC2514-2M, Vision Dimension) and a narrow near-infrared bandpass filter (BN850-25.4, Midwest Optical Systems). An LED positioned in front of the fly provided red light (622 nm) for CsChrimson activation at about 0.1 mW/mm^2^. Fly behaviour was recorded in 5-s trials. The optogenetic stimulus began at 2 s and lasted 1 s. The setup was controlled with custom Matlab code.

Fly behavior was classified manually per frame as resting, walking, grooming, flying, or other. To automatically detect the swing phases of the legs during walking in the side-view videos, we trained a deep neural network (DeepLabCut^75^) to automatically track all ipsilateral leg joints and other key points on the body (head-antenna joint, wing hinge, scutellum, posterior end of abdomen). The swing phase detection was based on two features: the vertical velocity of the tarsus and the velocity of a 2D vector pointing from the body-coxa joint to the tip of the tarsus. For each leg pair (front, middle, hind), we trained two long short-term memory (LSTM) networks to identify swing onsets or swing offsets based on these features (six networks in total). Each network used a bidirectional LSTM layer with 60 hidden units. The input data for each time point was a chunk of 20 samples (66.67 ms) of the two features around that time point. We labeled about 100 sequences per network, 90% of which were used for training and 10% of which were used for testing. Classification was done in Matlab (Deep Learning Toolbox). Where necessary, we manually corrected the first swing onset of each leg following the optogenetic stimulation (for the analysis shown in Fig. 1j,k), and we corrected missing or wrongly detected swing phases. For the analysis shown in Fig. 1k, legs that moved within 1 frame (3.33 ms) of each other were considered to move simultaneously. Tethered flies occasionally pushed the treadmill. Trials in which pushing occurred directly before or after the optogenetic activation were excluded from the analysis.

### Electrophysiology

Flies were prepared and recorded as described previously^62^. In brief, cold-anesthetized flies were tethered to a pyramid-shaped holder at their head using ultraviolet-curing glue. The proboscis was glued to minimize brain movement. To gain access to the brain, we opened the posterior side of the head and removed the overlying trachea, fat bodies, and ocellar ganglion. Flies were then positioned on top of the spherical treadmill. Collagenase was applied to a small region around the left or right RRN soma to rupture the overlaying neural sheath using a blunt patch-clamp electrode. Cells were recorded at 20 kHz (pCLAMP 11, Molecular Devices) using patch-clamp electrodes (7-12 MΩ) filled with intracellular saline (40 mM potassium aspartate, 10 mM HEPES, 1 mM EGTA, 4 mM MgATP, 0.5 mM Na3 GTP, 1 mM KCl, adjusted to 260–275 mOsm, pH 7.3). The treadmill ball was tracked at 50 Hz as described above. To immobilize the treadmill, we manually switched off the air flow. Side-view videos of the fly were recorded at 150 Hz using a camera (Basler acA1300-200um, Basler AG) equipped with a 75-mm lens (Tamron M112FM75, Vision Dimension). In flight experiments, the treadmill was lowered and wing movements were recorded with a wing beat tachometer^62^. Flight was induced by gentle air puffs. Data were recorded with Clampex 11 (Molecular Devices).

Action potentials were detected based on shape and amplitude (peakfinder function, Matlab). Thresholds were determined manually for each recording. The instantaneous spike rate was estimated by convolving the spike train with a Gaussian kernel (σ = 150 ms) and multiplying by the sampling rate. Forward, side, and angular treadmill velocities were low-pass filtered using a moving average filter with a width of 300 ms. Translational speed was calculated as the sum of the absolute smoothed forward and side velocity. Angular speed was calculated as the absolute smoothed angular velocity.

To create a binary time series indicating when the fly was walking, we first set all time points as walking where the translational speed or angular speed were larger than 0.3 mm/s or 10 deg/s, respectively. Then, epochs of up to 500 ms between detected walking events were also set to walking. Finally, we applied a hysteresis filter, which assigned epochs of up to 800 ms to the previous epoch (walking or non-walking). We ran the hysteresis filter twice, once forward in time and once backward. Pushing was annotated manually based on the side-view videos.

To create a binary time series indicating when the fly was flying, we first detected wingbeats by thresholding the tachometer signal. Similar to the walking detection, we then assigned short epochs of up to 10 ms between detected wingbeats as flight, and then applied a hysteresis filter to assign epochs of up to 100 ms to the previous epoch (flight or non-flight).

### Chemogenetic experiments

For chemogenetic activation experiments, we expressed the ATP receptor P2X_2_ (ref.^48^) together with GFP in RRNs. Control flies expressed GFP in RRNs, but lacked the P2X_2_ receptor. Flies were prepared as in the electrophysiology experiments. Collagenase was applied to a small region around the left or right RRN soma to rupture the overlaying neural sheath using a blunt patch-clamp electrode. Then, an electrode filled with 10 mM ATP (Sigma A9187) and 50 µM Alexa 488 (Invitrogen A10436) in extracellular saline was positioned close to the exposed soma. In each trial, ATP was applied for 500 ms with a pressure of 8-10 psi (PDES-02DX, NPI Electronic). Trials in which the animal was moving during or 2 s prior to the ATP injection were excluded from the analysis. At the end of each experiment, Alexa 488 diffusion was monitored in test injections under fluorescence illumination. If the pipette was clogged, the preceding trials were discarded. Without local desheathing, ATP application had no effect on behavior, suggesting that ATP mainly activated the exposed soma, not the soma on the other side of the brain. Treadmill velocities were low-pass filtered using a moving average filter with a width of 300 ms.

### Gap-crossing assay

The gap-crossing setup consisted of two 4-mm wide platforms spaced 3.5 mm apart inside a water-filled arena^51^. The platforms were illuminated from above by white light and four infrared LEDs (825 nm). Three additional LEDs provided red light (625 nm) for CsChrimson activation. Fly behavior was recorded at 150 Hz with a side-view camera (Basler acA1300-200um, Basler AG) equipped with a 50-mm lens (XR-400, SainSonic) and a near-infrared lowpass filter (LP780-25.4, Midwest Optical Systems). The setup was controlled with custom Matlab code.

Cold-anesthetized flies with clipped wings were transferred to one of the platforms. Then, we recorded multiple trials per animal in which we manually triggered a 1-s optogenetic stimulus. We determined post hoc whether the stimulus occurred in one of four phases. In Phase 0, flies stood in front of the gap. In Phase 1, at least one front leg had moved into the gap, while middle and hind legs were still on the platform. In Phase 2, at least one front leg had reached the other side of the gap, while middle and hind legs were still on the platform or had started moving to the other side. In Phase 3, all legs had moved to the other side of the gap or flies started climbing out of the gap, with the abdomen still in the gap. We manually annotated five behaviors after the optogenetic stimulation: no movement (resting), climbing, walking away from the gap, falling, and walking backward. We excluded trials in which flies climbed entirely on the side of the platforms and one trial in which a fly jumped. Moreover, we excluded trials for Phase 0 in which flies moved at stimulus onset.

The cross duration was calculated as the time between the first front leg moving toward the opposite side of the gap and all legs reaching the other side. The duration was calculated for trials in which the optogenetic stimulation was present when the first front leg started moving toward the other side, which included trials from Phase 0 and Phase 1.

Body lengths were determined as the distance between the neck and the end of the abdomen in video frames in which the abdomen and thorax formed a straight line, which was typically the case mid-crossing.

Latency to move was calculated as the time from stimulus onset to first forward (RRN) or first backward (MDN) movement of the body. For RRN, we only included data from Phase 0. For MDN, we included data from Phase 0 and Phase 1, because the onset of backward movement was clearly identifiable even when flies were already moving.

### Connectome analyses

#### Connectivity analyses in FAFB/FlyWire

RRNs were identified first in the hemibrain connectome^76^ by tracing the neurons in 3D in a template-registered confocal image stack of the split-GAL4 line and overlaying the skeletons with skeletons from the connectome (VVD Viewer; https://zenodo.org/records/14872850). RRNs were then identified in the FAFB/FlyWire connectome^26,38^ based on their main presynaptic neurons, which were annotated in both connectomes. RRN is cell-typed as CB0257 (FAFB/FlyWire connectome and Brain and Nerve Cord (BANC) connectome^29^) or CL249 (hemibrain connectome and male CNS connectome^30^).

The FAFB/FlyWire connectome and its synapse predictions^77^, neurotransmitter predictions^39^, and annotations^38^ were downloaded from flywire.ai or queried via CAVEclient^78^ and FAFBseg^25,26,38^. Connections between neurons with fewer than five synapses were excluded from connectivity analyses.

To analyze the shared output connectivity of RRN and BPN, we analyzed all 1- and 2-hop connections from each RRN and BPN to DNs. 2-hop connections with RRNs, BPNs, or DNs as intermediate neurons were excluded. Shared output connectivity was calculated for all RRNs or BPNs as the summed fraction of output synapses onto shared DNs, shared intermediate neurons, and intermediate neurons that project to shared DNs. To analyze the shared connectivity expected by chance, we shuffled the output connections from each RRN and BPN, using *n* = 10,000 shuffles^79^.

When analyzing the output connectivity of DNs in the FAFB/FlyWire brain connectome (Fig. 2f), we expressed the number of output synapses relative to the total number of output synapses that DNs make in the brain and nerve cord. To estimate this ratio, we queried the synapse locations of all DNs in the BANC connectome^29^, which spans the brain and nerve cord. Synapses were excluded for connections between neurons with fewer than five synapses. In BANC, the median fraction of DN outputs in the brain was 0.25, which we multiplied with the DN outputs in the FAFB/FlyWire connectome.

To analyze the net excitatory effect of RRN and BPN onto individual DNs across two hops, we retrieved the all-to-all connectivity matrix for all neurons of the RRN-BPN circuit and normalized all synapse weights to input fractions (sum of inputs to each postsynaptic neuron equals 1). Synapse weights from GABAergic and glutamatergic intermediate neurons were given negative values, reflecting their frequent inhibitory effect in *Drosophila*^80^. Connections from intermediate neurons with an unreliable neurotransmitter prediction (score <0.62, see ref.^39^) were set to zero (that is, excluded). Then, we multiplied the matrix with itself and added it to the original matrix, yielding a 2-hop effective connectivity strength^46,81^ between RRN or BPN and each DN, with positive values indicating net excitation^82^. The top DNs predicted to receive net excitation were the same when we assumed glutamatergic intermediate neurons to have an excitatory effect rather than an inhibitory effect.

To analyze the inputs of RRN and BPN from different brain neuropils, we first calculated the relative number of input synapses of each presynaptic neuron in those neuropils. The relative synapse counts were then multiplied with the number of synapses between each presynaptic neuron and RRN or BPN. In the public FAFB/FlyWire dataset used for our analysis (v783), one of the RRNs is missing its contralateral branch and a fraction of its input synapses. To mitigate effects on our presynaptic connectivity analysis, we pooled synapses from left and right neuropils and summed the weighted input synapses for both RRNs. We added the missing branch in the production dataset of the connectome for future studies.

#### Connectivity analyses in MANC

The connectivity of DNs of interest in the VNC was analyzed in the Male Adult Nerve Cord (MANC) connectome^28,43^. Connections between neurons with fewer than five synapses were excluded. Most DNs of interest in MANC were annotated by ref.^44^ or cross-referenced in BANC^29^. We identified two additional types (DNpe020, DNp52) in MANC based on their anatomy in BANC. One type (DNge138) could not be confidently identified in the VNC and was excluded from the analysis.

### Statistical analysis

Statistical analyses were done in Python and Matlab. To compare two groups of paired data, we used the Wilcoxon signed-rank test. To compare two groups of unpaired data, we used the Mann-Whitney U test. To compare multiple groups, we used a one-way ANOVA followed by Tukey’s HSD test. In the experiments analyzed with an ANOVA, the different groups contained the same animals, but animal identity could not be tracked across groups. Therefore, we treated the animals in each group as independent samples. In the figures, *, **, and *** indicate p < 0.05, p < 0.01, and p < 0.001, respectively. Test statistics are reported in the main text.

## Data availability

Behavioral and electrophysiology data generated in this study will be available for download at the time of publication. FAFB/FlyWire connectome data were analyzed from version 783, which was downloaded from https://codex.flywire.ai/api/download/ in June 2025. We used tables Connections Princeton No Threshold, Classification, and Neurotransmitter Type Predictions. MANC connectome data were analyzed from version 1.2.1, which was downloaded from https://neuprint.janelia.org/ in June 2025 using the neuPrint Python package.

## Code availability

Analyses were done in Matlab 2023a and Python 3.9. Code to recreate the figures is available at GitHub (https://github.com/chrisjdallmann/roadrunner/). Python code made use of CAVEclient, FAFBseg, neuPrint, and banc to query connectome data, NetworkX to analyze connectivity, NAVis, flybrains, and MeshParty to visualize neurons, scikit-learn for kernel density estimation, SciPy for statistics and time series filtering, and Matplotlib, seaborn, NumPy, and pandas for general computation and data visualization.

## Acknowledgements

We thank Hannah Volk for help with optogenetic experiments and Michael Dübbert, Sebastian Eberhardt, and Thomas Walter for technical support. We thank Ansgar Büschges, M. Eugenia Chiappe, Stephan Dietrich, and John C. Tuthill and his laboratory for feedback on the manuscript. We thank the BANC community for proofreading and annotating descending neurons and for providing an open resource that enabled early access to the connectome. We used stocks obtained from the Bloomington *Drosophila* Stock Center (NIH P40OD018537). This project received funding from the European Union’s Horizon Europe research and innovation program under the Marie Skłodowska-Curie grant agreement No. 101107596 to C.J.D. and the German Research Foundation (DFG) through the Emmy Noether program (DFG AC 371/1-1) and the Next Generation Networks for Neuroscience (NeuroNex) program (DFG AC 371/2-1) to J.M.A.

## Author contributions

C.J.D. and J.M.A. conceived the study. J.G. and K.I. generated the genetic driver line for RRNs. F.M.I. conducted optogenetic experiments of freely walking flies. F.M.I., F.C.M., and C.J.D. analyzed optogenetic experiments of freely walking flies. H.S. and C.J.D. conducted optogenetic experiments of tethered flies. C.J.D. analyzed optogenetic experiments of tethered flies. S. Liebscher conducted patch-clamp recordings. S. Liebscher and C.J.D. analyzed patch-clamp recordings. M.E. conducted and analyzed P2X_2_ experiments. F.M.I. conducted SPARC experiments. F.M.I, F.C.M., and C.J.D. analyzed SPARC experiments. J.G. and K.I. identified RRNs in the connectome. C.J.D. performed connectome analyses. E.L.S. and S. Liessem conducted gap-crossing experiments. E.L.S., S. Liessem, and C.J.D. analyzed gap-crossing experiments. C.J.D. visualized the results. C.J.D. and J.M.A. acquired funding. C.J.D. and J.M.A. wrote the manuscript with input from other authors. J.M.A. initiated and supervised the project.

## Competing interests

The authors declare no competing interests.

---

**Supplementary Fig. 1:**
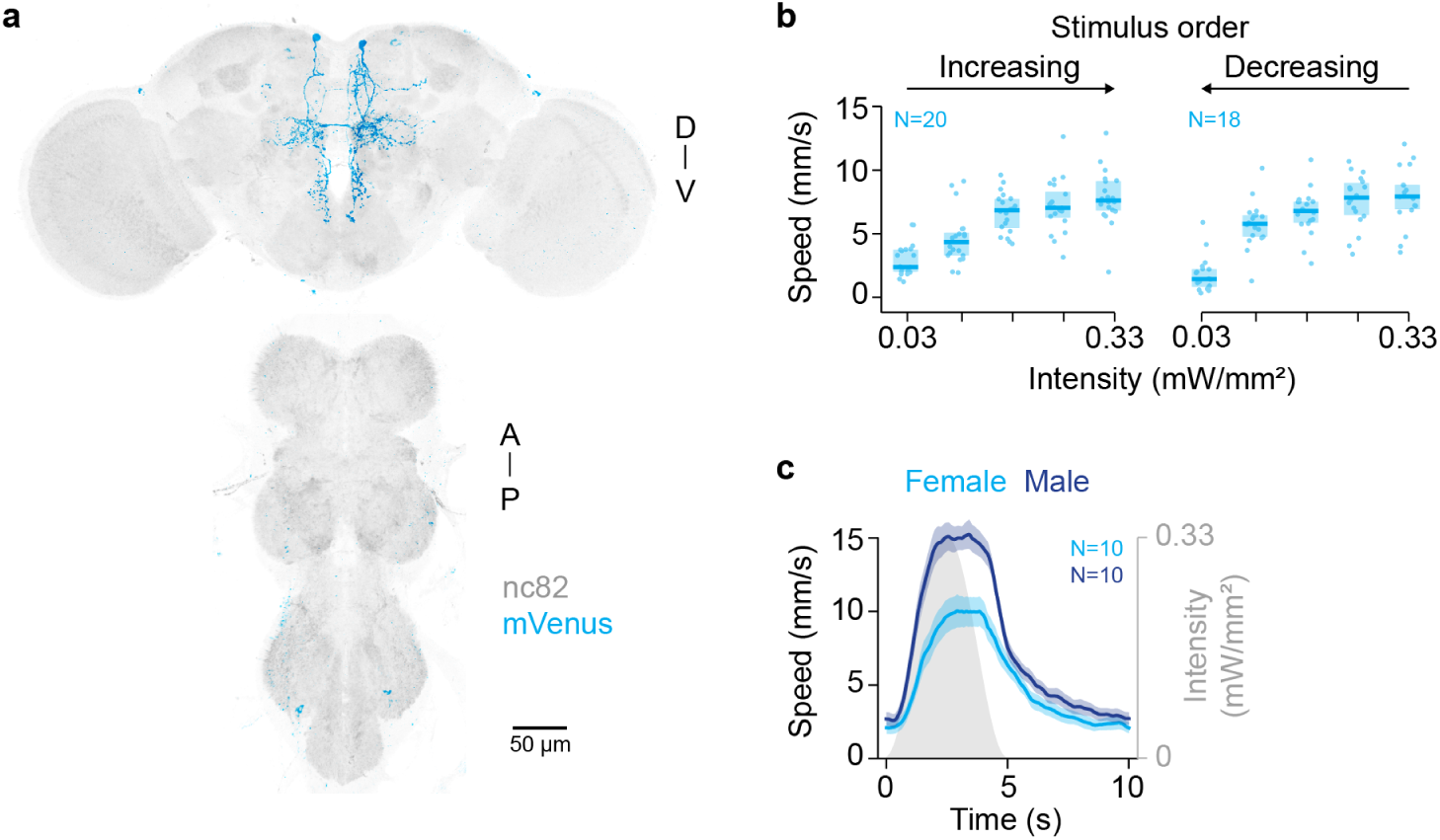
Anatomy and optogenetic activation of RRN. **a,** Light-microscopy image of the genetic driver line targeting the two RRNs in the brain. A, anterior; D, dorsal; P, posterior; V, ventral. **b,** Translational speeds of RRN>CsChrimson flies during optogenetic activation at different intensities. The intensity was either progressively increased (left) or decreased (right) over the course of the experiment. Dots, animal means; boxes, interquartile range with median. *N* = 20/18 flies (increasing/ decreasing); *n* = 9 activations per fly. **c,** Translational speed (mean ± s.e.m.) of female (light blue) and male (dark blue) RRN>CsChrimson flies during sinusoidal optogenetic activation (gray). *N* = 10 flies each; *n* = 40-50 activations per fly.

**Supplementary Fig. 2:**
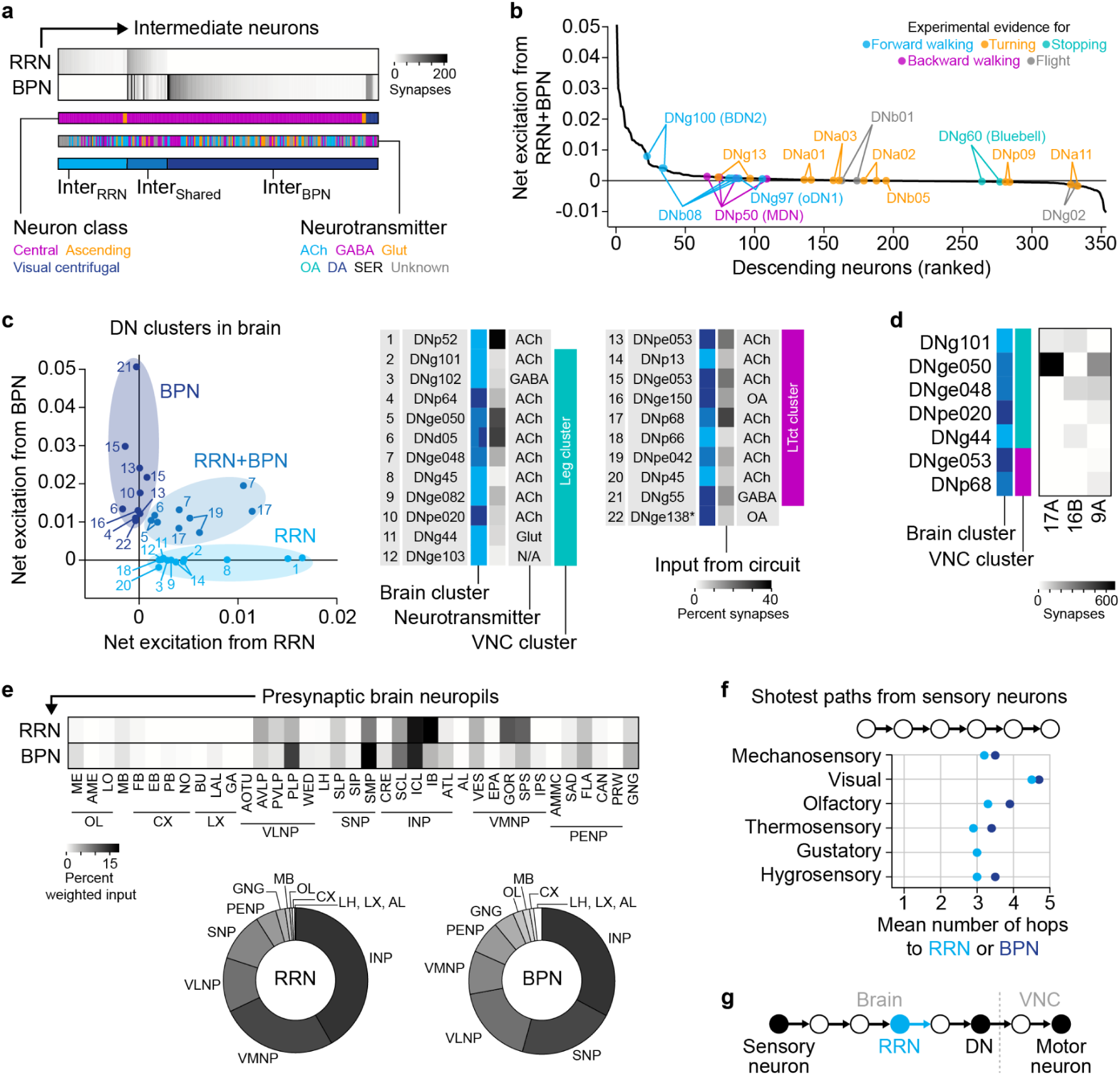
Classification and connectivity of neurons in the forward-walking circuit. **a,** Intermediate neurons postsynaptic to RRN, BPN, or both (FAFB/FlyWire connectome). ACh, acetylcholine; GABA, gamma-aminobutyric acid; Glut, glutamate; OA, octopamine; DA, dopamine, SER, serotonin; Unknown, neurons with a neurotransmitter prediction score smaller than 0.62 (ref.^39^). **b,** DNs ranked based on the predicted combined net excitation they receive from RRN and BPN. Colored dots indicate DNs previously implicated in locomotion control. **c,** Identities of DNs from Fig. 2g. Numbers, rows of connectivity heatmap in Fig. 2h; brain cluster, clusters on the left; neurotransmitter, neurotransmitter prediction (FAFB/FlyWire connectome); VNC cluster, clusters in Fig. 2h; grayscale column, percent input synapses from neurons of the RRN-BPN circuit; asterisk, DN type that could not be identified in the VNC. **d,** Connectivity of specific DNs downstream of RRN and BPN with walking-related interneurons in the VNC. 17A and 16B (E1 and I1 in ref.^41^), putative central-pattern-generating neurons for walking; 9A (chief 9A in ref.^47^), neurons involved in gating leg proprioception. Brain and VNC clusters as in **c**. Synapses were summed per DN type and interneuron type. **e,** Top, Weighted synaptic input from different brain neuropils onto RRN and BPN (Methods). Bottom, Weighted synaptic input from neuropils summed across broader brain regions. For neuropil and region names, see ref.^42^. **f,** Shortest paths from sensory neurons on the head to RRN or BPN. Dots, mean shortest path length between a sensory neuron of a modality and any RRN or BPN. **g,** Schematic of RRN being roughly equidistant to sensory and motor layers.

**Supplementary Fig. 3:**
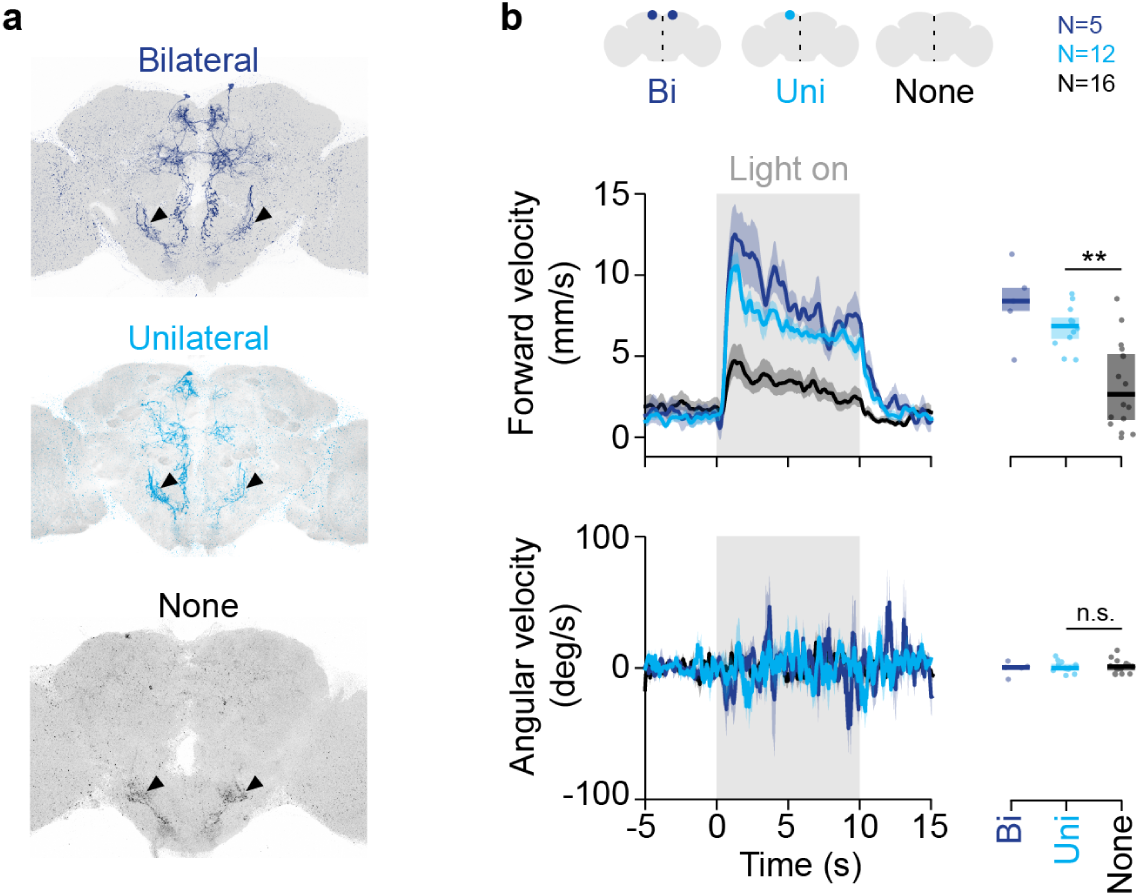
Stochastic labeling and optogenetic activation of RRN. **a,** Example expression patterns of RRNs using SPARC. Either both (bilateral), one (unilateral), or none of the neurons were labeled. Arrowheads indicate consistent off-target labeling. **b,** Forward and angular velocity (mean ± s.e.m.) of freely walking flies expressing CsChrimson in both (Bi), one (Uni), or none of the RRNs during optogenetic activation (gray; intensity of 0.33 mW/mm^2^). Positive angular velocity corresponds to ipsiversive turning. Dots, animal means (mean from 0 to 10 s); boxes, interquartile range with median. *N* = 5/12/16 flies (Bi/Uni/None); *n* = 9 activations per fly.

**Supplementary Fig. 4:**
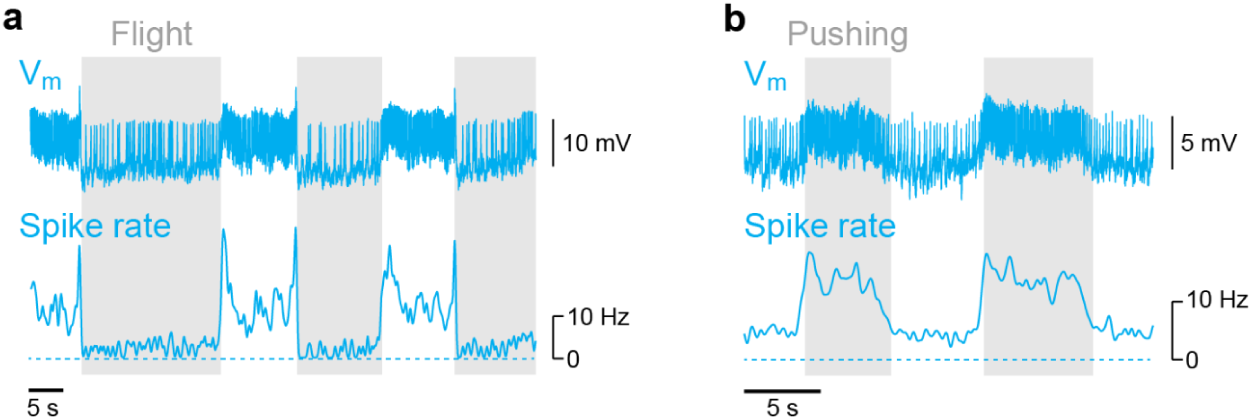
RRN activity during flight and pushing. **a,** Example patch-clamp recording of RRN in a tethered flying fly. **b,** Example patch-clamp recording of RRN in a tethered fly attempting to walk (pushing).

**Supplementary Fig. 5:**
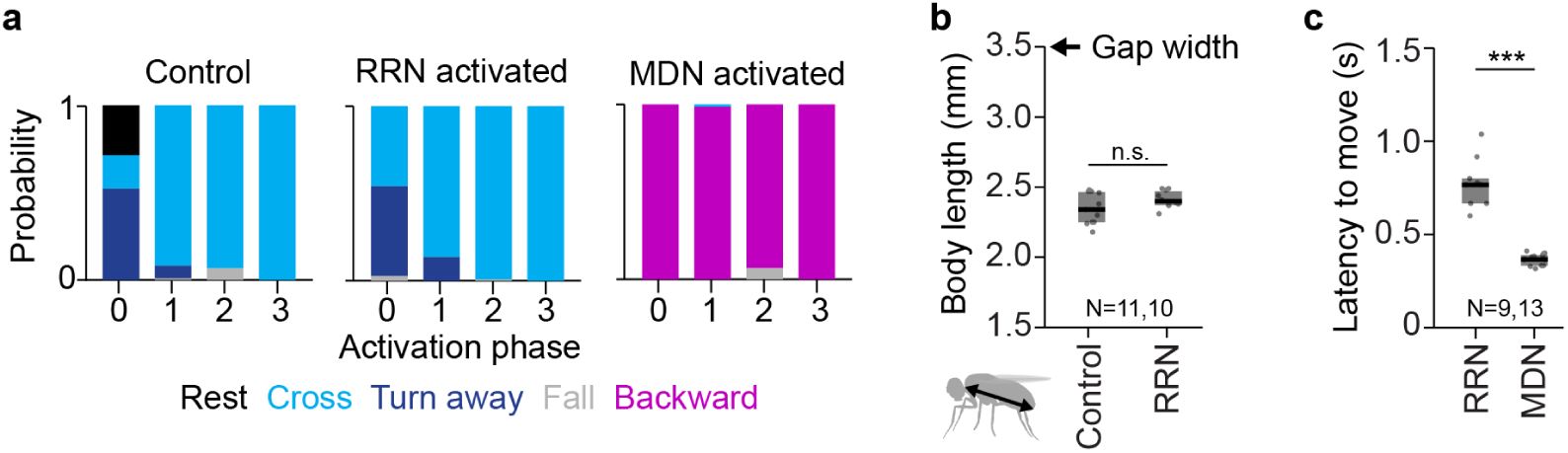
Behavior probability, body lengths, and latency to move in gap-crossing assay. **a,** Probability of behaviors of empty>CsChrimson (control), RRN>CsChrimson, and MDN>CsChrimson flies following optogenetic activation before (phase 0) or during (phases 1-3) gap-crossing. Behaviors pooled across flies. Same data as Fig. 5c but colored based on behavior. **b,** Body lengths of empty>CsChrimson (control) and RRN>CsChrimson flies. Body length corresponds to the distance between the neck and abdomen (inset). Dots, animal means; boxes, interquartile range with median. *N* = 11/10 (control/RRN). **c,** Latency to move of RRN>CsChrimson and MDN>CsChrimson flies in response to optogenetic stimulation. *N* = 9/13 flies (RRN/MDN); *n* = 1-16 movements per fly.

## Notes

### Competing Interest Statement

The authors have declared no competing interest.

## References

1. Leiras, R., Cregg, J. M. & Kiehn, O. Brainstem circuits for locomotion. Annu. Rev. Neurosci. 45, 63–85 (2022).

2. Arber, S. & Costa, R. M. Networking brainstem and basal ganglia circuits for movement. Nat. Rev. Neurosci. 23, 342–360 (2022).

3. Grillner, S. & El Manira, A. Current principles of motor control, with special reference to vertebrate locomotion. Physiol. Rev. 100, 271–320 (2020).

4. Lacroix-Ouellette, P. & Dubuc, R. Brainstem neural mechanisms controlling locomotion with special reference to basal vertebrates. Front. Neural Circuits 17, 910207 (2023).

5. Büschges, A. & Ache, J. M. Motor control on the move: from insights in insects to general mechanisms. Physiol. Rev. 105, 975–1031 (2025).

6. Caggiano, V., Leiras, R., Goñi-Erro, H., Masini, D., Bellardita, C., Bouvier, J., Caldeira, V., Fisone, G. & Kiehn, O. Midbrain circuits that set locomotor speed and gait selection. Nature 553, 455–460 (2018).

7. Josset, N., Roussel, M., Lemieux, M., Lafrance-Zoubga, D., Rastqar, A. & Bretzner, F. Distinct contributions of mesencephalic locomotor region nuclei to locomotor control in the freely behaving mouse. Curr. Biol. 28, 884–901.e3 (2018).

8. Roseberry, T. K., Lee, A. M., Lalive, A. L., Wilbrecht, L., Bonci, A. & Kreitzer, A. C. Cell-type-specific control of brainstem locomotor circuits by basal ganglia. Cell 164, 526–537 (2016).

9. Cregg, J. M., Sidhu, S. K., Leiras, R. & Kiehn, O. Basal ganglia–spinal cord pathway that commands locomotor gait asymmetries in mice. Nat. Neurosci. 27, 716–727 (2024).

10. Carbo-Tano, M., Lapoix, M., Jia, X., Thouvenin, O., Pascucci, M., Auclair, F., Quan, F. B., Albadri, S., Aguda, V., Farouj, Y., Hillman, E. M. C., Portugues, R., Del Bene, F., Thiele, T. R., Dubuc, R. & Wyart, C. The mesencephalic locomotor region recruits V2a reticulospinal neurons to drive forward locomotion in larval zebrafish. Nat. Neurosci. 26, 1775–1790 (2023).

11. Berg, E. M., Mrowka, L., Bertuzzi, M., Madrid, D., Picton, L. D. & El Manira, A. Brainstem circuits encoding start, speed, and duration of swimming in adult zebrafish. Neuron 111, 372–386.e4 (2023).

12. Sapkal, N., Mancini, N., Kumar, D. S., Spiller, N., Murakami, K., Vitelli, G., Bargeron, B., Maier, K., Eichler, K., Jefferis, G. S. X. E., Shiu, P. K., Sterne, G. R. & Bidaye, S. S. Neural circuit mechanisms underlying context-specific halting in *Drosophila*. Nature 634, 191–200 (2024).

13. Feng, K., Khan, M., Minegishi, R., Müller, A., Van De Poll, M. N., Swinderen, B. V. & Dickson, B. J. A central steering circuit in *Drosophila*. bioRxiv 2024.06.27.601106 (2024).

14. Capelli, P., Pivetta, C., Soledad Esposito, M. & Arber, S. Locomotor speed control circuits in the caudal brainstem. Nature 551, 373–377 (2017).

15. Cregg, J. M., Leiras, R., Montalant, A., Wanken, P., Wickersham, I. R. & Kiehn, O. Brainstem neurons that command mammalian locomotor asymmetries. Nat. Neurosci. 23, 730–740 (2020).

16. Wyart, C., Jia, X., Kumar, A., Peysson, N., Carbo-Tano, M., Fidelin, K., Didelot, M., Lejeune, C. & Grosse-Wentrup, M. All-optical investigation reveals a hierarchical organization of vsx2+ reticulospinal neurons coordinating steering and forward locomotion. Res. Sq. 10.21203/rs.3.rs-7216842/v1 (2025).

17. Lau, J. Y. N., Fitzgerald, J. E. & Bianco, I. H. Supraspinal commands have a modular organization that is behavioral context specific. Curr. Biol. 35, 4408–4425.e6 (2025).

18. Bidaye, S. S., Laturney, M., Chang, A. K., Liu, Y., Bockemühl, T., Büschges, A. & Scott, K. Two brain pathways initiate distinct forward walking programs in *Drosophila*. Neuron 108, 469–485.e8 (2020).

19. Namiki, S., Ros, I. G., Morrow, C., Rowell, W. J., Card, G. M., Korff, W. & Dickinson, M. H. A population of descending neurons that regulates the flight motor of *Drosophila*. Curr. Biol. 32, 1189–1196.e6 (2022).

20. Rayshubskiy, A., Holtz, S. L., Bates, A., Vanderbeck, Q. X., Capdevila, L. S. & Wilson, R. I. Neural circuit mechanisms for steering control in walking *Drosophila*. eLife 13, RP102230 (2024).

21. Yang, H. H., Brezovec, B. E., Serratosa Capdevila, L., Vanderbeck, Q. X., Adachi, A., Mann, R. S. & Wilson, R. I. Fine-grained descending control of steering in walking *Drosophila*. Cell 187, 6290–6308.e27 (2024).

22. Mussells Pires, P., Zhang, L., Parache, V., Abbott, L. F. & Maimon, G. Converting an allocentric goal into an egocentric steering signal. Nature 626, 808–818 (2024).

23. Westeinde, E. A., Kellogg, E., Dawson, P. M., Lu, J., Hamburg, L., Midler, B., Druckmann, S. & Wilson, R. I. Transforming a head direction signal into a goal-oriented steering command. Nature 626, 819–826 (2024).

24. Ros, I. G., Omoto, J. J. & Dickinson, M. H. Descending control and regulation of spontaneous flight turns in *Drosophila*. Curr. Biol. 34, 531–540.e5 (2024).

25. Zheng, Z., Lauritzen, J. S., Perlman, E., Robinson, C. G., Nichols, M., Milkie, D., Torrens, O., Price, J., Fisher, C. B., Sharifi, N., Calle-Schuler, S. A., Kmecova, L., Ali, I. J., Karsh, B., Trautman, E. T., Bogovic, J. A., Hanslovsky, P., Jefferis, G. S. X. E., Kazhdan, M., Khairy, K., et al. A complete electron microscopy volume of the brain of adult *Drosophila melanogaster*. Cell 174, 730–743.e22 (2018).

26. Dorkenwald, S., Matsliah, A., Sterling, A. R., Schlegel, P., Yu, S., McKellar, C. E., Lin, A., Costa, M., Eichler, K., Yin, Y., Silversmith, W., Schneider-Mizell, C., Jordan, C. S., Brittain, D., Halageri, A., Kuehner, K., Ogedengbe, O., Morey, R., Gager, J., Kruk, K., et al. Neuronal wiring diagram of an adult brain. Nature 634, 124–138 (2024).

27. Azevedo, A., Lesser, E., Phelps, J. S., Mark, B., Elabbady, L., Kuroda, S., Sustar, A., Moussa, A., Khandelwal, A., Dallmann, C. J., Agrawal, S., Lee, S.-Y. J., Pratt, B., Cook, A., Skutt-Kakaria, K., Gerhard, S., Lu, R., Kemnitz, N., Lee, K., Halageri, A., et al. Connectomic reconstruction of a female *Drosophila* ventral nerve cord. Nature 631, 360–368 (2024).

28. Takemura, S., Hayworth, K. J., Huang, G. B., Januszewski, M., Lu, Z., Marin, E. C., Preibisch, S., Xu, C. S., Bogovic, J., Champion, A. S., Cheong, H. S., Costa, M., Eichler, K., Katz, W., Knecht, C., Li, F., Morris, B. J., Ordish, C., Rivlin, P. K., Schlegel, P., et al. A connectome of the male *Drosophila* ventral nerve cord. eLife 13, RP97769 (2024).

29. Bates, A. S., Phelps, J. S., Kim, M., Yang, H. H., Matsliah, A., Ajabi, Z., Perlman, E., Delgado, K. M., Osman, M. A. M., Salmon, C. K., Gager, J., Silverman, B., Renauld, S., Collie, M. F., Fan, J., Pacheco, D. A., Zhao, Y., Patel, J., Zhang, W., Serratosa Capdevilla, L., et al. Distributed control circuits across a brain-and-cord connectome. bioRxiv 2025.07.31.667571 (2025).

30. Berg, S., Beckett, I. R., Costa, M., Schlegel, P., Januszewski, M., Marin, E. C., Nern, A., Preibisch, S., Qiu, W., Takemura, S., Fragniere, A. M. C., Champion, A. S., Adjavon, D.-Y., Cook, M., Gkantia, M., Hayworth, K. J., Huang, G. B., Kampf, F., Katz, W. T., Lu, Z., et al. Sexual dimorphism in the complete connectome of the *Drosophila* male central nervous system. bioRxiv 2025.10.09.680999 (2025).

31. Meissner, G. W., Nern, A., Dorman, Z., DePasquale, G. M., Forster, K., Gibney, T., Hausenfluck, J. H., He, Y., Iyer, N. A., Jeter, J. &, et al. A searchable image resource of *Drosophila* GAL4 driver expression patterns with single neuron resolution. eLife 12, e80660 (2023).

32. Isaacman-Beck, J., Paik, K. C., Wienecke, C. F. R., Yang, H. H., Fisher, Y. E., Wang, I. E., Ishida, I. G., Maimon, G., Wilson, R. I. & Clandinin, T. R. SPARC enables genetic manipulation of precise proportions of cells. Nat. Neurosci. 23, 1168–1175 (2020).

33. Brezovec, B. E., Berger, A. B., Hao, Y. A., Chen, F., Druckmann, S. & Clandinin, T. R. Mapping the neural dynamics of locomotion across the *Drosophila* brain. Curr. Biol. 34, 710–726.e4 (2024).

34. Klapoetke, N. C., Murata, Y., Kim, S. S., Pulver, S. R., Birdsey-Benson, A., Cho, Y. K., Morimoto, T. K., Chuong, A. S., Carpenter, E. J., Tian, Z., Wang, J., Xie, Y., Yan, Z., Zhang, Y., Chow, B. Y., Surek, B., Melkonian, M., Jayaraman, V., Constantine-Paton, M., Wong, G. K.-S., et al. Independent optical excitation of distinct neural populations. Nat. Methods 11, 338–346 (2014).

35. Mohammad, F., Stewart, J. C., Ott, S., Chlebikova, K., Chua, J. Y., Koh, T.-W., Ho, J. & Claridge-Chang, A. Optogenetic inhibition of behavior with anion channelrhodopsins. Nat. Methods 14, 271–274 (2017).

36. Govorunova, E. G., Sineshchekov, O. A., Janz, R., Liu, X. & Spudich, J. L. Natural light-gated anion channels: A family of microbial rhodopsins for advanced optogenetics. Science 349, 647–650 (2015).

37. Bisen, R. S., Iqbal, F. M., Cascino-Milani, F., Bockemühl, T. & Ache, J. M. Nutritional state-dependent modulation of insulin-producing cells in *Drosophila*. eLife 13, RP98514 (2025).

38. Schlegel, P., Yin, Y., Bates, A. S., Dorkenwald, S., Eichler, K., Brooks, P., Han, D. S., Gkantia, M., dos Santos, M., Munnelly, E. J., Badalamente, G., Serratosa Capdevila, L., Sane, V. A., Fragniere, A. M. C., Kiassat, L., Pleijzier, M. W., Stürner, T., Tamimi, I. F. M., Dunne, C. R., Salgarella, I., et al. Whole-brain annotation and multi-connectome cell typing of *Drosophila*. Nature 634, 139–152 (2024).

39. Eckstein, N., Bates, A. S., Champion, A., Du, M., Yin, Y., Schlegel, P., Lu, A. K.-Y., Rymer, T., Finley-May, S., Paterson, T., Parekh, R., Dorkenwald, S., Matsliah, A., Yu, S.-C., McKellar, C., Sterling, A., Eichler, K., Costa, M., Seung, S., Murthy, M., et al. Neurotransmitter classification from electron microscopy images at synaptic sites in Drosophila melanogaster. Cell 187, 2574–2594.e23 (2024).

40. Bidaye, S. S., Machacek, C., Wu, Y. & Dickson, B. J. Neuronal control of *Drosophila* walking direction. Science 344, 97–101 (2014).

41. Pugliese, S. M., Chou, G. M., Abe, E. T. T., Turcu, D., Lancaster, J. K., Tuthill, J. C. & Brunton, B. W. Connectome simulations identify a central pattern generator circuit for fly walking. bioRxiv 2025.09.12.675944 (2025).

42. Ito, K., Shinomiya, K., Ito, M., Armstrong, J. D., Boyan, G., Hartenstein, V., Harzsch, S., Heisenberg, M., Homberg, U., Jenett, A., Keshishian, H., Restifo, L. L., Rössler, W., Simpson, J. H., Strausfeld, N. J., Strauss, R. & Vosshall, L. B. A systematic nomenclature for the insect brain. Neuron 81, 755–765 (2014).

43. Marin, E. C., Morris, B. J., Stuerner, T., Champion, A. S., Krzeminski, D., Badalamente, G., Gkantia, M., Dunne, C. R., Eichler, K., Takemura, S., Tamimi, I. F. M., Fang, S., Moon, S. S., Cheong, H. S. J., Li, F., Schlegel, P., Berg, S., FlyEM Project Team, Card, G. M., Costa, M., et al. Systematic annotation of a complete adult male *Drosophila* nerve cord connectome reveals principles of functional organisation. eLife 13, RP97766 (2024).

44. Stürner, T., Brooks, P., Capdevila, L. S., Morris, B. J., Javier, A., Fang, S., Gkantia, M., Cachero, S., Beckett, I. R., Champion, A. S., Moitra, I., Richards, A., Klemm, F., Kugel, L., Namiki, S., Cheong, H. S. J., Kovalyak, J., Tenshaw, E., Parekh, R., Schlegel, P., et al. Comparative connectomics of the descending and ascending neurons of the *Drosophila* nervous system: stereotypy and sexual dimorphism. Nature 643, 158–172 (2025).

45. Namiki, S., Dickinson, M. H., Wong, A. M., Korff, W. & Card, G. M. The functional organization of descending sensory-motor pathways in *Drosophila*. eLife 7, e34272 (2018).

46. Cheong, H. S., Eichler, K., Stürner, T., Asinof, S. K., Champion, A. S., Marin, E. C., Oram, T. B., Sumathipala, M., Venkatasubramanian, L., Namiki, S., Siwanowicz, I., Costa, M., Berg, S., Janelia FlyEM Project Team, Jefferis, G. S. & Card, G. M. Transforming descending input into motor output: An analysis of the *Drosophila* Male Adult Nerve Cord connectome. eLife 13, RP96084 (2025).

47. Dallmann, C. J., Luo, Y., Agrawal, S., Mamiya, A., Chou, G. M., Cook, A., Sustar, A., Brunton, B. W. & Tuthill, J. C. Selective presynaptic inhibition of leg proprioception in behaving *Drosophila*. Nature 647, 445–453 (2025).

48. Lima, S. Q. & Miesenböck, G. Remote control of behavior through genetically targeted photostimulation of neurons. Cell 121, 141–152 (2005).

49. Fujiwara, T., Brotas, M. & Chiappe, M. E. Walking strides direct rapid and flexible recruitment of visual circuits for course control in *Drosophila*. Neuron 110, 2124–2138.e8 (2022).

50. Chen, C.-L., Aymanns, F., Minegishi, R., Matsuda, V. D. V., Talabot, N., Günel, S., Dickson, B. J. & Ramdya, P. Ascending neurons convey behavioral state to integrative sensory and action selection brain regions. Nat. Neurosci. 26, 682–695 (2023).

51. Pick, S. & Strauss, R. Goal-driven behavioral adaptations in gap-climbing *Drosophila*. Curr. Biol. 15, 1473–1478 (2005).

52. Dahlhoff, S., Liessem, S., Iqbal, F. M., Palacios-Muñoz, A., Cascino-Milani, F., Erginkaya, M., Diniz, A. M., Gorostiza, E. A., Büschges, A., Clemens, J. & Ache, J. M. Control of walking direction by descending and dopaminergic neurons in *Drosophila*. bioRxiv 2025.07.22.666129 (2025).

53. Shik, M., Severin, F. & Orlovskiĭ, G. Control of walking and running by means of electric stimulation of the midbrain. Biofizika 11, 659–666 (1966).

54. Sirota, M. G., Di Prisco, G. V. & Dubuc, R. Stimulation of the mesencephalic locomotor region elicits controlled swimming in semi-intact lampreys. Eur. J. Neurosci. 12, 4081–4092 (2000).

55. Musienko, P. E., Zelenin, P. V., Lyalka, V. F., Gerasimenko, Y. P., Orlovsky, G. N. & Deliagina, T. G. Spinal and supraspinal control of the direction of stepping during locomotion. J. Neurosci. 32, 17442–17453 (2012).

56. Schwenkgrub, J., Harrell, E. R., Bathellier, B. & Bouvier, J. Deep imaging in the brainstem reveals functional heterogeneity in V2a neurons controlling locomotion. Sci. Adv. 6, eabc6309 (2020).

57. Kaplan, H. S., Nichols, A. L. A. & Zimmer, M. Sensorimotor integration in *Caenorhabditis elegans* : a reappraisal towards dynamic and distributed computations. Philos. Trans. R. Soc. B Biol. Sci. 373, 20170371 (2018).

58. Tammero, L. F. & Dickinson, M. H. Collision-avoidance and landing responses are mediated by separate pathways in the fruit fly, *Drosophila melanogaster*. J. Exp. Biol. 205, 2785–2798 (2002).

59. Berthé, R. & Lehmann, F.-O. Body appendages fine-tune posture and moments in freely manoeuvring fruit flies. J. Exp. Biol. jeb.122408 (2015).

60. Ache, J. M., Namiki, S., Lee, A., Branson, K. & Card, G. M. State-dependent decoupling of sensory and motor circuits underlies behavioral flexibility in *Drosophila*. Nat. Neurosci. 22, 1132–1139 (2019).

61. Liessem, S., Asinof, S. K., Nern, A., Sumathipala, M., Rogers, E., Erginkaya, M., Dallmann, C. J., Card, G. M. & Ache, J. M. Parallel neuronal ensembles control behavior across sensorimotor levels in Drosophila. bioRxiv 2025.12.13.693955 (2025).

62. Liessem, S., Held, M., Bisen, R. S., Haberkern, H., Lacin, H., Bockemühl, T. & Ache, J. M. Behavioral state-dependent modulation of insulin-producing cells in *Drosophila*. Curr. Biol. 33, 449–463.e5 (2023).

63. Cheong, H. S. J., Boone, K. N., Bennett, M. M., Salman, F., Ralston, J. D., Hatch, K., Allen, R. F., Phelps, A. M., Cook, A. P., Phelps, J. S., Erginkaya, M., Lee, W.-C. A., Card, G. M., Daly, K. C. & Dacks, A. M. Organization of an ascending circuit that conveys flight motor state in *Drosophila*. Curr. Biol. 34, 1059–1075.e5 (2024).

64. Seeds, A. M., Ravbar, P., Chung, P., Hampel, S., Midgley, F. M., Mensh, B. D. & Simpson, J. H. A suppression hierarchy among competing motor programs drives sequential grooming in *Drosophila*. eLife 3, e02951 (2014).

65. Hampel, S., Franconville, R., Simpson, J. H. & Seeds, A. M. A neural command circuit for grooming movement control. eLife 4, e08758 (2015).

66. Özdil, P. G., Arreguit, J., Scherrer, C., Ijspeert, A. & Ramdya, P. Centralized brain networks underlie body part coordination during grooming. bioRxiv 2024.12.17.628844 (2024).

67. Syed, D. S., Ravbar, P. & Simpson, J. H. Inhibitory circuits control leg movements during *Drosophila* grooming. eLife 14, RP106446 (2025).

68. Triphan, T., Poeck, B., Neuser, K. & Strauss, R. Visual targeting of motor actions in climbing *Drosophila*. Curr. Biol. 20, 663–668 (2010).

69. Flash, T. & Hochner, B. Motor primitives in vertebrates and invertebrates. Curr. Opin. Neurobiol. 15, 660–666 (2005).

70. Braun, J., Hurtak, F., Wang-Chen, S. & Ramdya, P. Descending networks transform command signals into population motor control. Nature 630, 686–694 (2024).

71. Xu, L., Zhu, B., Zhu, Z., Tao, X., Zhang, T., El Manira, A. & Song, J. Separate brainstem circuits for fast steering and slow exploratory turns. Nat. Commun. 16, 3207 (2025).

72. Chockley, A. S., Dinges, G. F., Di Cristina, G., Ratican, S., Bockemühl, T. & Büschges, A. Subsets of leg proprioceptors influence leg kinematics but not interleg coordination in *Drosophila melanogaster* walking. J. Exp. Biol. 225, jeb244245 (2022).

73. Walter, T. & Couzin, I. D. TRex, a fast multi-animal tracking system with markerless identification, and 2D estimation of posture and visual fields. eLife 10, e64000 (2021).

74. Schindelin, J., Arganda-Carreras, I., Frise, E., Kaynig, V., Longair, M., Pietzsch, T., Preibisch, S., Rueden, C., Saalfeld, S., Schmid, B., Tinevez, J.-Y., White, D. J., Hartenstein, V., Eliceiri, K., Tomancak, P. & Cardona, A. Fiji: An open-source platform for biological-image analysis. Nat. Methods 9, 676–682 (2012).

75. Mathis, A., Mamidanna, P., Cury, K. M., Abe, T., Murthy, V. N., Mathis, M. W. & Bethge, M. DeepLabCut: Markerless tracking of user-defined features with deep learning. Nat. Neurosci. 21, 1281–1289 (2018).

76. Scheffer, L. K., Xu, C. S., Januszewski, M., Lu, Z., Takemura, S., Hayworth, K. J., Huang, G. B., Shinomiya, K., Maitlin-Shepard, J., Berg, S., Clements, J., Hubbard, P. M., Katz, W. T., Umayam, L., Zhao, T., Ackerman, D., Blakely, T., Bogovic, J., Dolafi, T., Kainmueller, D., et al. A connectome and analysis of the adult *Drosophila* central brain. eLife 9, e57443 (2020).

77. Yu, S.-C., Bae, J. A., Matsliah, A., Dorkenwald, S., Gager, J., Hebditch, J., Silverman, B., Willie, K. P., Willie, R., Burke, A. T., Macrina, T., Seung, S. & Murthy, M. New synapse detection in the whole-brain connectome of *Drosophila*. bioRxiv 2025.07.11.664377 (2025).

78. Dorkenwald, S., Schneider-Mizell, C. M., Brittain, D., Halageri, A., Jordan, C., Kemnitz, N., Castro, M. A., Silversmith, W., Maitin-Shephard, J., Troidl, J., Pfister, H., Gillet, V., Xenes, D., Bae, J. A., Bodor, A. L., Buchanan, J., Bumbarger, D. J., Elabbady, L., Jia, Z., Kapner, D., et al. CAVE: Connectome Annotation Versioning Engine. Nat. Methods 22, 1112–1120 (2025).

79. Lesser, E., Azevedo, A. W., Phelps, J. S., Elabbady, L., Cook, A., Syed, D. S., Mark, B., Kuroda, S., Sustar, A., Moussa, A., Dallmann, C. J., Agrawal, S., Lee, S.-Y. J., Pratt, B., Skutt-Kakaria, K., Gerhard, S., Lu, R., Kemnitz, N., Lee, K., Halageri, A., et al. Synaptic architecture of leg and wing premotor control networks in *Drosophila*. Nature 631, 369–377 (2024).

80. Liu, W. W. & Wilson, R. I. Glutamate is an inhibitory neurotransmitter in the *Drosophila* olfactory system. Proc. Natl. Acad. Sci. 110, 10294–10299 (2013).

81. Li, F., Lindsey, J. W., Marin, E. C., Otto, N., Dreher, M., Dempsey, G., Stark, I., Bates, A. S., Pleijzier, M. W., Schlegel, P., Nern, A., Takemura, S., Eckstein, N., Yang, T., Francis, A., Braun, A., Parekh, R., Costa, M., Scheffer, L. K., Aso, Y., et al. The connectome of the adult *Drosophila* mushroom body provides insights into function. eLife 9, e62576 (2020).

82. Lee, S.-Y. J., Dallmann, C. J., Cook, A., Tuthill, J. C. & Agrawal, S. Divergent neural circuits for proprioceptive and exteroceptive sensing of the *Drosophila* leg. Nat. Commun. 16, 4105 (2025).

